# The Food Additive Xanthan Gum Drives Adaptation of the Human Gut Microbiota

**DOI:** 10.1101/2021.06.02.446819

**Authors:** Matthew P. Ostrowski, Sabina Leanti La Rosa, Benoit J. Kunath, Andrew Robertson, Gabriel Pereira, Live H. Hagen, Neha J. Varghese, Ling Qiu, Tianming Yao, Gabrielle Flint, James Li, Sean McDonald, Duna Buttner, Nicholas A. Pudlo, Matthew K. Schnizlein, Vincent B. Young, Harry Brumer, Thomas Schmidt, Nicolas Terrapon, Vincent Lombard, Bernard Henrissat, Bruce Hamaker, Emiley A Eloe-Fadrosh, Ashootosh Tripathi, Phillip B. Pope, Eric Martens

## Abstract

The diets of industrialized countries reflect the increasing use of processed foods, often with the introduction of novel food additives. Xanthan gum is a complex polysaccharide with unique rheological properties that have established its use as a widespread stabilizer and thickening agent^1^. However, little is known about its direct interaction with the gut microbiota, which plays a central role in digestion of other, chemically-distinct dietary fiber polysaccharides. Here, we show that the ability to digest xanthan gum is surprisingly common in industrialized human gut microbiomes and appears to be contingent on the activity of a single bacterium that is a member of an uncultured bacterial genus in the family *Ruminococcaceae*. We used a combination of enrichment culture, multi-omics, and recombinant enzyme studies to identify and characterize a complete pathway in this uncultured bacterium for the degradation of xanthan gum. Our data reveal that this keystone degrader cleaves the xanthan gum backbone with a novel glycoside hydrolase family 5 (GH5) enzyme before processing the released oligosaccharides using additional enzymes. Surprisingly, some individuals harbor a *Bacteroides* species that is capable of consuming oligosaccharide products generated by the keystone *Ruminococcaceae* or a purified form of the GH5 enzyme. This *Bacteroides* symbiont is equipped with its own distinct enzymatic pathway to cross-feed on xanthan gum breakdown products, which still harbor the native linkage complexity in xanthan gum, but it cannot directly degrade the high molecular weight polymer. Thus, the introduction of a common food additive into the human diet in the past 50 years has promoted the establishment of a food chain involving at least two members of different phyla of gut bacteria.

## Introduction

Evidence is accumulating that food additives impact the symbiosis between humans and their associated gut microbiomes, in some cases promoting intestinal inflammation and metabolic syndrome^2^ or promoting certain pathogens^3^. Often used as thickeners and emulsifiers, polysaccharides are a prominent subset of these food additives. Since dietary polysaccharides other than starch typically transit the upper intestinal tract undigested, polysaccharide-based additives can potentially exert their influence by altering the composition and function of the microbiome, which can in turn impact host health^4,5^. Given generally regarded as safe (GRAS) approval by the United States Food and Drug Administration in 1968, xanthan gum (XG) is an exopolysaccharide produced by the bacterium *Xanthamonas campestris* that has been increasingly used in the food supply for the last 50 years. This polymer has the same β−1,4-linked backbone as cellulose, but contains trisaccharide branches on alternating glucose residues consisting of α-1,3-mannose, β-1,2-glucuronic acid, and terminal β-1,4-mannose (**Figure 1a**). The terminal β-D-mannose and the inner α-D-mannose are variably pyruvylated at the 4,6-position or acetylated at the 6-position, respectively, with amounts determined by specific *X. campestris* strains and culture conditions^6^. XG is typically added at concentrations of 0.05-0.5% to foods including bakery products, condiments, and ice cream^7^. XG is also used as a replacement for gluten in a gluten-free diet, which is a vital component for limiting intestinal inflammation in patients with celiac disease, a lifelong condition estimated to affect 0.7-1.4% of the population and increasing in prevalence^8^. In gluten-free baked goods, XG can be consumed in up to gram quantities per serving. Although small doses of XG have not been connected to immediate health impacts, its fate in the digestive tract is unknown^9^. The low-level, but constant consumption of XG by a large portion of the population in the industrialized world and its higher intake by specific subpopulations highlight the need to understand the effects of this polysaccharide food additive on the ecology of the human gut microbiota.

**Figure 1.**
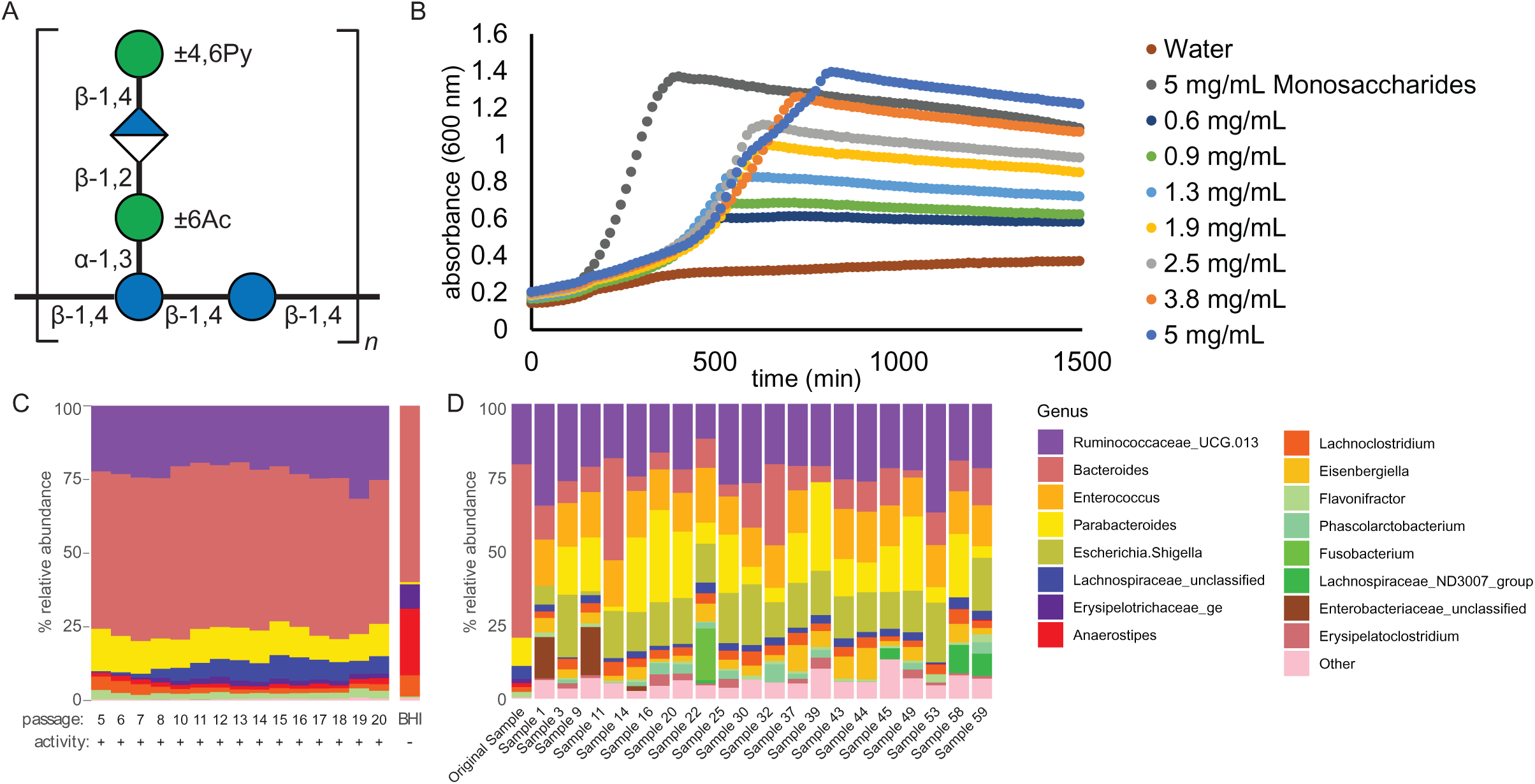
R. UCG13 was a common factor across xanthan gum degrading cultures. **a**, Xanthan gum is a repeating structure of glucose (blue circles), mannose (green circles), and glucuronic acid (blue and white diamond). The inner and outer mannose residues are variably modified by acetylation and pyruvylation, respectively. **b**, Growth curves of the original xanthan-degrading culture showed greater culture density as xanthan gum concentration was increased (n=12, SEM ≤ 3%), and, **c**, displayed relatively stable composition over sequential passaging. Passaging the culture on BHI-blood plates resulted in a loss of R. UCG13 as well as xanthan degrading activity. **d**, An additional 20 samples were sequentially passaged in xanthan containing media (10x) and analyzed for composition by 16S rRNA sequencing (16 of the most abundant genus are displayed for clarity). All cultures shared an abundant OTU, classified as R. UCG13.

### A member of an uncultured bacterial genus is a keystone XG degrader in the human gut microbiome

To identify potential XG degrading bacteria in the human gut microbiome, we surveyed a group of healthy 18-20 year-old adults using a bacterial enrichment culture strategy in which a partially defined minimal medium was combined with XG as the main carbon source^10^. This medium was inoculated directly with feces that had been collected in anaerobic preservation buffer within 24 h prior to testing and cultures containing individual samples were passaged 3 times with 1-2 days of growth in between. Growth of the final passage was monitored quantitatively for bacterial growth or for loss of the gel-like viscosity that is characteristic of XG in solution. We originally found one XG-degrading culture (see *Materials and Methods*) and in an expanded experiment in which 60 individuals were sampled we identified 30 positive cultures, indicating that the ability of intestinal bacteria to degrade XG is unexpectedly common among the population surveyed.

Experiments with a culture derived from one positive subject revealed that bacterial growth depended on the amount of XG provided in the medium, demonstrating specificity for this nutrient (**Figure 1b**). Attempts to enrich the causal XG-consuming organism(s) with additional passaging (total of 10-20 times) consistently yielded stable mixed microbial cultures that contained multiple operational taxonomic units (OTUs; between 12-22 OTUs per culture with relative abundance ≥0.5%) (**Figure 1c, 1d**). While these cultures had commonalities at the genus level, there was surprisingly only one OTU that was ≥0.5% and common across all 21 enrichment cultures examined. This common OTU was identified as a member of Ruminococcaceae uncultured genus 13 (R. UCG13) in the Silva database^11^ (**Figure 1d, Supplemental Table 1**). Plating and passaging the same culture used in **Figure 1b** on a commonly used anaerobic solid medium (brain-heart infusion with 10% horse blood) resulted in loss of two previously abundant Gram-positive OTUs (loss defined as <0.01% relative abundance), which included the R. UCG13 OTU and corresponded with loss of the XG-degrading phenotype when plate-passaged bacteria were re-inoculated into medium with XG (**Figure 1c**).

Despite R. UCG13 and a *Bacteroides* OTU being present at >20% relative abundance in the original 12 OTU community, we repeatedly failed to isolate pure cultures that could degrade XG using different solid media that are effective for Gram-positive and -negative bacteria (the abundant *Bacteroides* OTU was captured multiple times, while R. UCG13 was never isolated). Dilution of the active 12-OTU community to extinction in medium supplemented with either XG or an equal amount of its component monosaccharides, resulted in loss of growth on XG at higher dilutions than simple sugars (**Extended Data 2**). This observation suggests that the ability to degrade XG in the medium conditions we employed requires multiple OTUs to be present (*i.e*., diluted into the same well together), which could be explained by either multiple species being directly involved or the necessity of other species to promote sufficient growth conditions for the XG degrader^12^. Collectively, these results suggest that a member of an uncultured Ruminococcaceae genus is necessary for XG degradation but may be unable to grow in isolation in our media formulation.

### Community sequencing identifies two putative XG utilization loci in R. UCG13 and *Bacteroides intestinalis*

To identify XG-degrading genes within our bacterial consortium, we performed combined metagenomics and metatranscriptomics analysis on the original XG-degrading culture, using samples harvested throughout growth in liquid medium with XG (**Extended Data 3**). From these samples, we reconstructed 18 metagenome assembled genomes (MAGs), 7 that were high quality (completion >90% and contamination <5%) (**Supplemental Table 2**). To connect 16S rRNA genes to MAGs, we performed additional long-read sequencing that yielded 2 MAGs that were complete circular chromosomes and one of these was identified as R. UCG13 (with four complete 16S rRNA operons, three of which were identical to the R. UCG13 OTU). This circular R. UCG13 genome was distantly related (47.26% Average Amino Acid Identity, AAI) to the recently cultured bacterium *Monoglobus pectinolyticus*^13^. Annotation of carbohydrate-active enzymes (CAZymes) in the R. UCG13 MAG revealed a single locus encoding several highly expressed enzymes that are candidates for XG degradation (**Figure 2, Extended Data 3**). These included a polysaccharide lyase family 8 (PL8) with distant homology to known xanthan lyases from soil bacteria *Paenibacillus alginolyticus* XL-1^14^ (36% identity/73% coverage) and *Bacillus* sp. GL1^15^ (32% identity/81% coverage; **Figure 2**). Xanthan lyases typically remove the terminal pyruvylated mannose prior to depolymerization of the β−1,4-glucose backbone, leaving a 4,5 unsaturated residue at the glucuronic acid position, although some tolerate non-pyruvylated mannose^16,17^. This same locus also contained two GH5 enzymes, a family that includes endoglucanases and xyloglucanases, with the potential to cleave the xanthan gum backbone^18^, a GH88 to remove the unsaturated glucuronic acid residue produced by the PL8^19^, and two GH38s which could potentially cleave the α-D-mannose^20^. Two carbohydrate esterases (CEs) could potentially remove the acetylation from the mannose^21^. Secretion signal prediction detected possible signal peptidase I (SPI) motifs for the two GH5s and one of the CEs (CE-A), while the other enzymes lacked detectable membrane localization and secretion signals^22^. In addition to putative enzymes to cleave the glycosidic bonds contained within xanthan gum, this locus also contained proteins predicted to be involved in sensing, binding, and transporting sugars and oligosaccharides.

**Figure 2.**
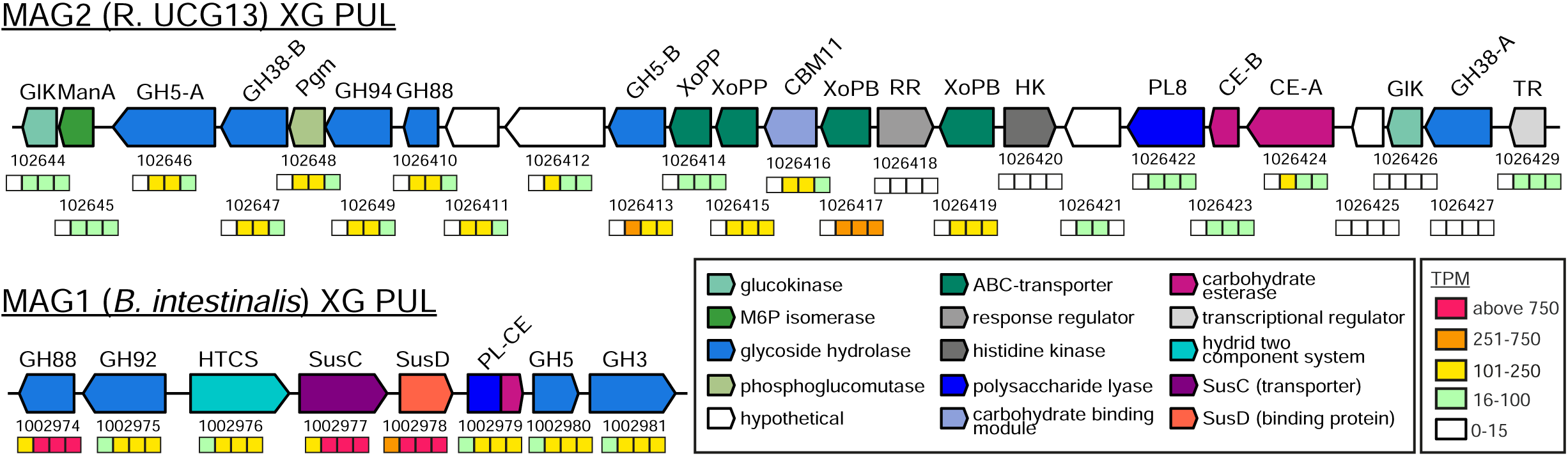
Metagenomics, metatranscriptomics, and activity-guided proteomics identified two putative xanthan gum degrading loci. Putative xanthan utilization loci color-coded and annotated by predicted protein family. The four boxes below each gene are colored to represent expression levels of each gene at timepoints taken throughout the culture’s growth on xanthan gum. MAG taxonomy is indicated in parentheticals.

Co-localization and expression of genes that saccharify a common polysaccharide into discrete polysaccharide utilization loci (PULs) is common in the Gram-negative *Bacteroidetes*^23^. Although not present in most XG-degrading cultures, we obtained a second circular MAG affiliated to *B. intestinalis,* which was conspicuously the most abundant OTU (up to ∼50%) in the mixed species culture that it was derived from (**Figure 1**). This MAG contained a putative PUL that was highly expressed during growth on XG (**Figure 2, Extended Data 3**) and encodes hallmark SusC-/SusD-like proteins, a sensor/regulator, and predicted GH88, GH92 and GH3 enzymes, which could potentially cleave the unsaturated β-glucuronyl, α-mannosyl, and β-glucosyl linkages in XG, respectively. Like the candidate gene cluster in R. UCG13, this PUL also contained a GH5 enzyme, which was assigned to subfamily GH5_5. Finally, a putative polysaccharide lyase (PL) was predicted, remotely related to alginate lyases^24,25^, as a candidate for removing the terminal mannose. In addition to the lyase domain, this multi-modular protein contains a carbohydrate esterase domain (CE) that could remove the acetyl groups positioned on the mannose. Extensive work has been conducted to characterize the substrate-specificity of PULs, which is demonstrated by hundreds of genomes with characterized and predicted PULs in the PUL database (PUL-DB)^26^. However, this database only harbored a single genome with a partially related homolog of the *B. intestinalis* PUL (*B. salyersiae* WAL 10018 PUL genes HMPREF1532_01924-HMPREF1532_01938), highlighting the diversity of polysaccharide utilization machinery that remains for discovery and characterization.

Interestingly, neutral monosaccharide analysis from our XG-degrading culture showed a relatively stable 1:1 ratio of glucose:mannose in the residual polysaccharide in the culture over time, implying that lyase-digested xanthan gum was not accumulating as growth progressed (**Extended Data 3**). This could be due to fast depolymerization and cellular importation of the XG polymer following lyase removal of the terminal mannose. Alternatively, the data are also consistent with a degradation model in which the XG backbone is cleaved prior to subsequent hydrolysis of the repeating pentasaccharide, a pathway that has not been characterized in XG-degrading bacteria and might result in pentasaccharides with full linkage complexity being transported into the bacterial cell before depolymerization^27,28^.

### R. UCG13 produces a unique endo-acting xanthanase activity

To investigate the cellular location of the enzymes responsible for xanthan degradation, the original 12-OTU culture was grown in XG medium and separated into filtered cell-free supernatant, cells that were washed to remove supernatant and resuspended or lysed, or lysed cells with supernatant. Incubation of these fractions with XG and subsequent analysis by thin layer chromatography (TLC) revealed that the cell-free supernatant was capable of depolymerizing XG into large oligosaccharides, while the intracellular fraction was required to further saccharify these products into smaller components (**Extended Data 4**). Liquid chromatography-mass spectrometry (LC-MS) analysis of cell-free supernatant incubated with XG revealed the presence of pentameric and decameric oligosaccharides matching the structure of xanthan gum (**Figure 3a**), supporting the model described above in which secreted *endo*-cleaving enzymes first hydrolyze native XG before xanthan lyase and other enzymes cleave the attached sidechains.

**Figure 3.**
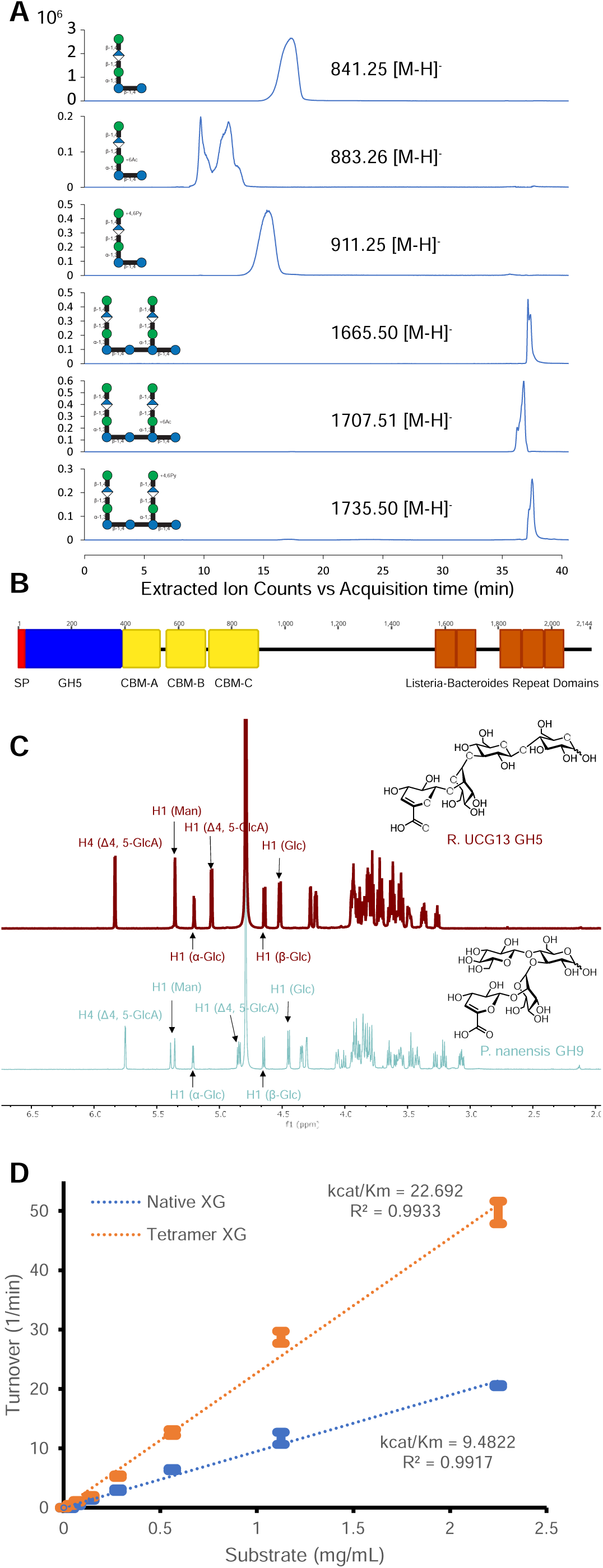
R. UCG13 encodes a novel GH5 that depolymerizes native xanthan gum. **a**, Extracted ion chromatograms showing various acetylated and pyruvylated penta- and deca-saccharides produced by incubating culture supernatant with XG. **b**, Annotated domains of the xanthan-degrading *Ru*GH5a, showing its signal peptide (SP), three carbohydrate binding modules (CBMs), and multiple Listeria-Bacteroides repeat domains. **c**, Proton NMR contrasting tetrasaccharide products obtained from incubating lyase-treated xanthan gum with either *Ru*GH5a or *P. nanensis* GH9. **d**, Kinetics of *Ru*GH5a on native and lyase-treated xanthan gum (error bars represent mean and standard deviation, n=4)

To identify enzymes responsible for XG hydrolysis and determine their cellular location, we grew three independent cultures in liquid medium containing XG and subjected cell-free supernatants to ammonium sulfate precipitation. Each of the resuspended protein preparations was able to hydrolyze XG as demonstrated by a complete loss of viscosity after overnight incubation. We proceeded to fractionate each sample with various purification methods (defined in methods), collecting and pooling xanthan-degrading fractions for subsequent purification steps and taking three different purification paths (**Extended Data 5**). Interestingly, the most pure sample obtained ran primarily as a large smear when loaded onto an SDS-PAGE gel, but separated into distinct bands after boiling, suggesting the formation of a multimeric protein complex, which is reminiscent of cellulosomes or other complexes^29^ (**Extended Data 5**). Proteomic analysis of the samples from the three different activity-guided fractionation experiments yielded 33 proteins present across all three experiments, including 22 from R. UCG13, 11 of which were annotated as CAZymes (**Extended Data 5, Supplemental Table 3**). While most of the proteins were either detected in low amounts or lacked functional predictions consistent with polysaccharide degradation, one of the most abundant proteins across all three samples was one of two GH5 enzymes (*Ru*GH5a) encoded in the previously identified R. UCG13 candidate xanthan locus.

*Ru*GH5a consists of an N-terminal signal peptide sequence, its main catalytic domain (which does not classify into any of the GH5 subfamilies), and 3 tandem carbohydrate-binding modules (CBMs), which are often associated with CAZymes and can facilitate polysaccharide degradation (**Figure 3b**)^30^. The protein also contains a significant portion of undefined sequence and *Listeria-Bacteroides* repeat domains (pfam^31^ PF09479), a β-grasp domain originally characterized from the invasion protein InlB used by *Listeria monocytogenes* for host cell entry^32,33^. These small repeat domains are thought to be involved in protein-protein interactions and are almost exclusively found in extracellular bacterial multidomain proteins. To test the activity of *Ru*GH5a on XG we expressed recombinant forms of the entire protein (*Ru*GH5a full), the GH5 domain only (*Ru*GH5a GH5-only), and the GH5 domain with either one (*Ru*GH5a GH5+CBM-A), two (*Ru*GH5a GH5+CBM-A/B), or all three of the CBMs (*Ru*GH5a GH5+CBMs, hereafter referred to as *Ru*GH5a for simplicity). All but the full-length construct yielded reasonably pure proteins, but only constructs with the GH5 and all three CBMs showed activity on xanthan gum, suggesting a critical role in catalysis for these CBMs. (**Extended Data 6**). The alternate GH5 (*Ru*GH5b) was also expressed in a variety of forms but did not display any activity on XG (**Extended Data 6)**.

Analysis of the reaction products showed that *Ru*GH5a releases pentasaccharide repeating units of XG, with various acetylation and pyruvylation (including di-acetylation as previously described^34^), and larger decasaccharide structures (**Figure 3a**). While isolation of homogenous pentameric oligosaccharides proved difficult, coincubation of XG with *Ru*GH5a and a *Bacillus sp.* PL8 facilitated the isolation of pure tetrasaccharide followed by in-depth 1D and 2D NMR structural characterization (**Extended Data 7**), which was useful in determining *Ru*GH5a regiospecificity in the XG backbone. Surprisingly, the NMR analysis suggested that *Ru*GH5a cleaves XG at the reducing end of the non-branching backbone glucosyl residue (**Figure 3c, Extended Data 7**). This contrasts with the product of other known xanthanases (such as the GH9 from *Paenibacillus nanensis*^35^ or the β-D-glucanase in *Bacillus* sp. strain GL1^15^), which hydrolyze xanthan at the reducing end of the branching glucose, demonstrating a hitherto unknown enzymatic mechanism for the degradation of XG. While *Ru*GH5a displayed little activity on other polysaccharides (**Extended Data 6**), it was able to hydrolyze both native and lyase-treated XG with comparable specificity, once more in contrast to most previously known xanthanases, which show ≥600-fold preference for the lyase-treated substrate^35^ (**Figure 3d**). One exception is the xanthanase from *Microbacterium* sp. XT11, which also cleaves native and lyase-treated xanthan gum with similar kinetic specificity^28^; however, this enzyme only produces intermediate XG fragments from the complete polysaccharide, whereas *Ru*GH5a can cleave XG down to its repeating pentasaccharide unit. Together these data highlight the novelty of *Ru*GH5a, which may be part of a multimeric protein complex *in vivo* and possesses a unique enzymatic mechanism and specificity.

### R. UCG13 encodes all the enzymes required for XG saccharification

In contrast to characterized PL8^14,17,36^ xanthan lyases, the R. UCG13 PL8 showed no activity on the complete XG polymer but removed the terminal mannose from xanthan pentasaccharides produced by *Ru*GH5a (**Extended Data 8**). This further supports the model in which the GH5 first depolymerizes XG, followed by further saccharification of the XG repeating unit, likely inside the cell. Both R. UCG13 carbohydrate esterases were able to remove acetyl groups from acetylated xanthan pentasaccharides (**Extended Data 8**). The tetrasaccharide produced by the PL8 was processed by the GH88 and both GH38s, which were able to saccharify the resulting trisaccharide (**Extended Data 8**). The GH94 catalyzed the phosphorolysis of cellobiose in phosphate buffer, completing the full saccharification of XG (**Extended Data 8**). Apparent redundancy of several enzymes (CEs and GH38s) could be partially explained by different cell location (e.g. CE-A has an SPI signal while CE-B does not), unique specificities for oligosaccharide variants in size or modification (i.e. acetylation or pyruvylation), additional polysaccharides that the locus targets, or evolutionary hypotheses where this locus is in the process of streamlining or expanding. Additional support for the involvement of this locus in XG degradation is provided by RNA-seq based whole genome transcriptome analysis, which showed the induction of genes in this cluster when the community was grown on XG compared to another polysaccharide (polygalacturonic acid, PGA) that also supports R. UCG13 abundance (**Extended Data 9**).

### *B. intestinalis* cross-feeds on XG oligosaccharides with its xanthan utilization PUL

Although R. UCG13 was recalcitrant to culturing efforts, we isolated several bacteria from the original consortium, including a representative strain of the *Bacteroides intestinalis* that was the most abundant (**Figure 1c**) and also harbors a highly expressed candidate PUL for XG degradation (**Figure 2**). While this strain was unable to grow on native XG as a substrate, we hypothesized that it may be equipped to utilize smaller XG fragments, such as those released during growth by R. UCG13 via its GH5 enzyme. Using the recombinant *Ru*GH5a, we generated sufficient quantities of mixed XG oligosaccharides (XGOs; primarily pentameric, but also some decameric oligosaccharides) to test growth of *Bacteroides intestinalis*. While isolates of *P. distasonis* and *B. clarus* from the same culture showed little or no growth (**Extended Data 9**), the *B. intestinalis* strain achieved comparable density on the XGOs as cultures grown on a stoichiometric mixture of the monosaccharides that compose XG, suggesting that it utilizes most or all of the sugars contained in the XGOs (**Figure 4a**). Consistent with the candidate *B. intestinalis* XGOs PUL being involved in this phenotype, all of the genes in this locus were activated >100-fold (and some >1000-fold) during growth on XGOs compared to a glucose-grown reference (**Figure 4b**). Whole genome RNA-seq analysis of the *B. intestinalis* strain grown on XGOs revealed that the identified PUL was the most highly upregulated in the genome, further validating its role in metabolism of XGOs (**Extended Data 9**). Interestingly, *Ru*GH5a XGOs treated with PL8 continued to support *B. intestinalis* growth, but tetramer generated from the *P. nanensis* GH9 and PL8 failed to support any growth (**Extended Data 9**). Growth was rescued in the presence of glucose but not in the presence of *Ru*GH5a XGOs to upregulate the PUL (**Extended Data 9**), suggesting that either the *B. intestinalis* transporters or enzymes are incapable of processing this isomeric substrate.

**Figure 4.**
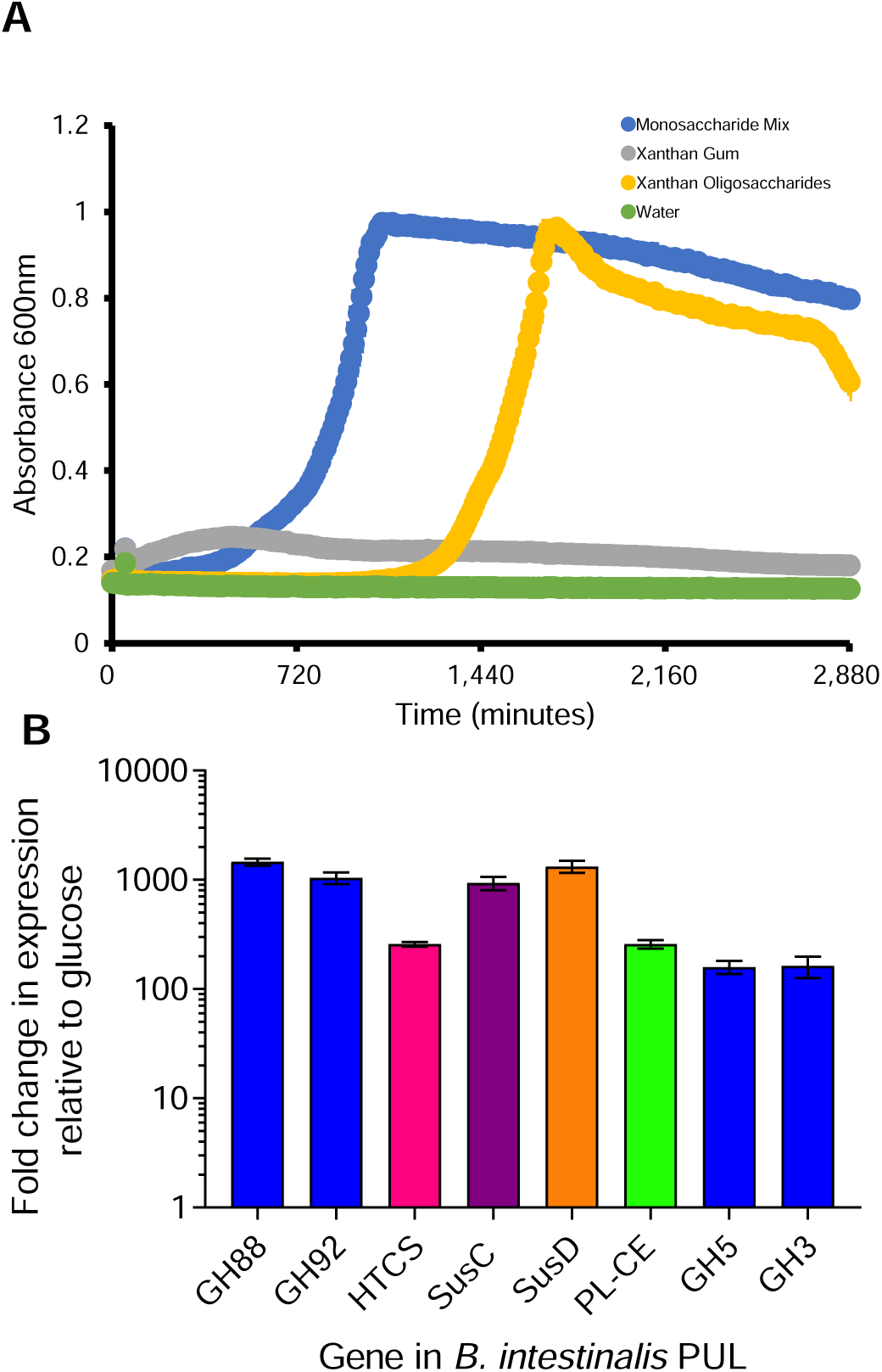
*B. intestinalis* cross-feeds on xanthan oligosaccharides. **a**, Growth curves of *B. intestinalis* isolated from the original xanthan-degrading culture. (curves represent mean and SEM, n=2). **b**, Fold-change in expression of *B. intestinalis* genes when grown on xanthan oligosaccharides relative to glucose.

To further test the role of the identified *B. intestinalis* PUL in XGOs degradation, we tested recombinant forms of its constituent enzymes for their ability to degrade XGOs, and confirmed the activity of several enzymes. The carbohydrate esterase domain C-terminal to the PL-CE bimodular protein was able to remove acetyl groups from acetylated xanthan pentasaccharides (**Extended Data 8**). While we were unable to detect xanthan lyase activity for the PL-CE enzyme on full length XG or oligosaccharides it is likely that this enzyme or another lyase acts to remove the terminal mannose residue since the GH88 was able to remove the corresponding 4,5 unsaturated glucuronic acid residue from the corresponding tetrasaccharide that would be generated by its action (**Extended Data 8**). The GH92 was active on the trisaccharide produced by the GH88 as observed by loss of the trisaccharide and formation of cellobiose (**Extended Data 8**). Finally, the GH3 was active on cellobiose, but did not show activity on either tri- or tetra-saccharide, suggesting that this enzyme may be the final step in *B. intestinalis* saccharification of XGOs (**Extended Data 8**). Signal peptidase II (SPII) secretion signals were predicted for the GH5, GH3, GH88, and SusD proteins while the GH92, PL-CE, HTCS, and SusC all had SPI motifs^22^. While signal peptides do not definitively determine cellular location, these predictions and accumulated knowledge of Sus-type systems in Bacteroidetes suggest a “selfish” model in which saccharification occurs primarily in the periplasm^23,37^ (**Extended Data 1**).

### Metagenomics suggests additional cross-feeding modalities

To determine if the original consortium was representative of all our XG-degrading cultures, we performed metagenomic sequencing on 20 additional XG-degrading communities and retrieved 16 high-quality and 3 low-quality R. UCG13 MAGs as well as an unbinned contig affiliated with R. UCG13 (**Supplemental Table 2**). We found that the R. UCG13 XG utilization locus is extremely well conserved across these cultures with only one variation in gene content, the insertion of a GH125 coding gene, and >95% amino acid identity (**Extended Data 10**). The additional GH125 gene was observed in most loci (14/17), suggesting that this gene provides a complementary, but non-essential function, possibly as an accessory α-mannosidase^38^. In contrast, only a subset of the samples (4/17) contained the *B. intestinalis* XGOs PUL, which showed essentially complete conservation in cultures that contained this PUL (**Extended Data 10**). Across all these cultures, R. UCG13 accounted for an average of only 23.1% ± 1.2 (SEM) of the total culture (**Figure 1d**), suggesting that additional microbes beyond *B. intestinalis* may cross-feed on products released by R. UCG13, either from degradation products of XG or by using other growth substrates generated by R. UCG13. For example, we found that the bacterial communities in samples 1, 22, and 59 contain other microbes belonging to the Bacteroidaceae family that harbor a PUL with a GH88, GH92, and GH3, suggesting that these bacteria can potentially metabolize XG-derived tetramers (**Extended Data 10**).

### Xanthan utilization loci are widespread in modern microbiomes

Next, we asked whether XG inclusion in the modern diet could have increased the prevalence of the R. UCG13 and *B. intestinalis* xanthan loci compared to other populations, such as hunter-gatherers, that are less likely to be exposed to this food additive. Using each locus as a query, we searched several publicly available fecal metagenome datasets collected from populations worldwide. All modern populations sampled displayed some presence of the R. UCG13 XG locus, with the Chinese and Japanese cohorts being the highest (up to 51% in one cohort) **(Figure 5**). The *B. intestinalis* locus was less prevalent, with two industrialized population datasets (Japan and Denmark/Spain) lacking any incidence. Where the locus was present, its prevalence ranged from 1-11%. The three hunter-gatherer or non-industrialized populations sampled, the Yanomami, Hadza, and Burkina Faso had no detected presence of either the R. UCG13 or *B. intestinalis* locus.

**Figure 5.**
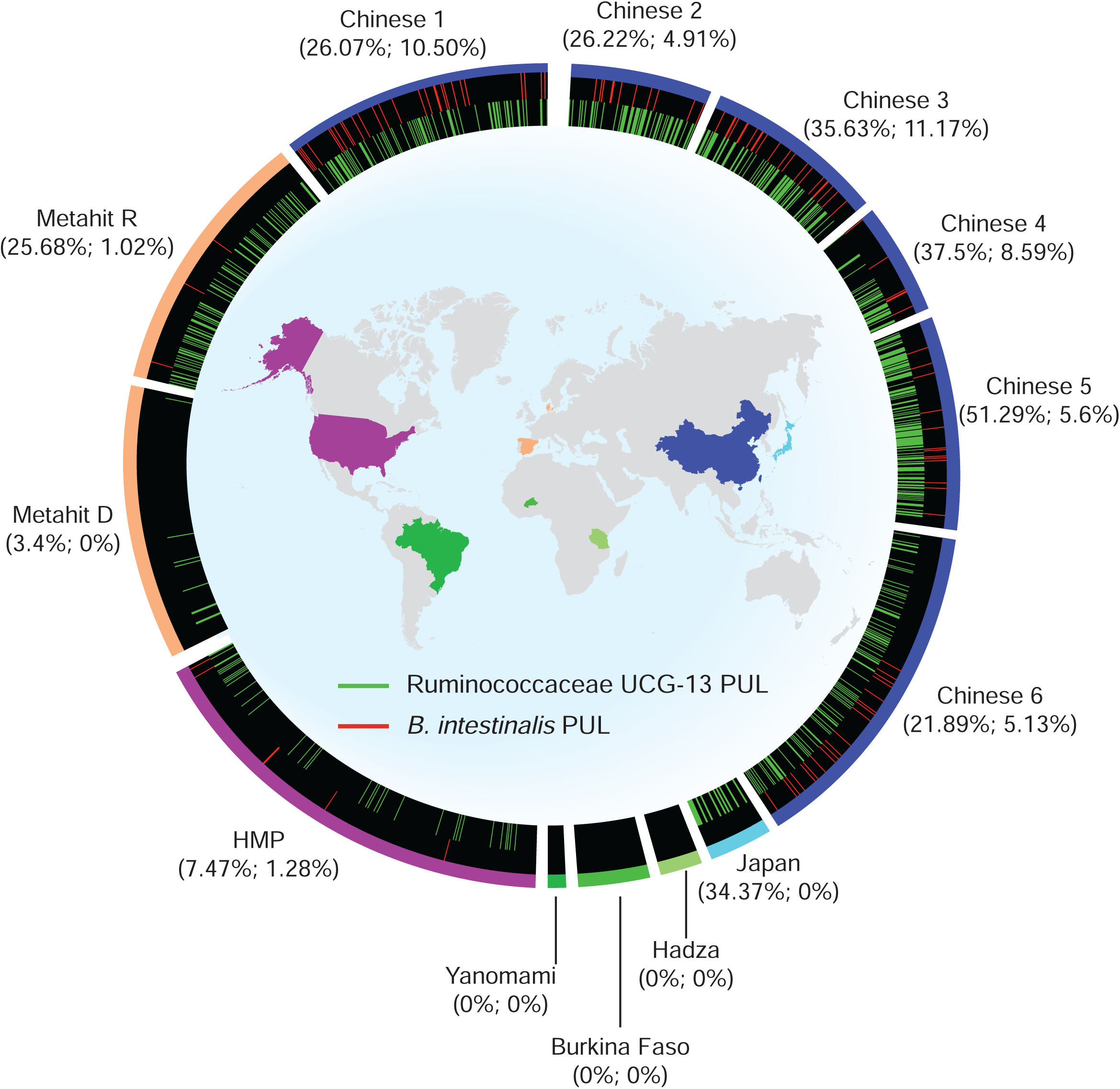
Xanthan degrading loci are present in modern human microbiomes but not in hunter-gatherers’. Multiple microbiome metagenome datasets were searched for the presence or absence of the R. UCG13 and *B. intestinalis* xanthan loci. Map colors correspond to where populations were sampled for each dataset displayed on the outside of the figure. Circle segments are sized proportionately to total number of individuals sampled for each dataset. Lines represent presence of either the R. UCG13 xanthan locus (green) or the *B. intestinalis* xanthan locus (red). Percentages display the total abundance of R. UCG13 or *B. intestinalis* locus in each dataset.

Although the size of the hunter-gatherer datasets is relatively small, excluding the possibility of a false negative suggests several equally intriguing hypotheses. Most obviously, inclusion of XG in the modern diet may have driven either the colonization or expansion of R. UCG13 (and to a lesser extent *B. intestinalis*) into the gut communities of numerous human populations. This is in concordance with the observations of Daly et al. who found that a set of volunteers fed xanthan gum for an extended period produced stool with increased probability and degree of xanthan degradation^39^. Alternatively, the modern microbiome is drastically different than that of hunter-gatherers and these differences simply correlate with the abundance of R. UCG13, rather than any causal effect of XG in the diet. Another possible hypothesis is that the microbiomes of hunter-gatherer populations can degrade XG, but through completely different microbes and pathways, a hypothesis that could be tested by culturing microbes from hunter-gatherer populations.

To further probe the presence of the identified XG utilization genes in other environments, we conducted an expanded LAST search^40^ of both loci in 72,491 sequenced bacterial isolates and 102,860 genome bins extracted from 13,415 public metagenomes, as well as 21,762 public metagenomes that are part of the Integrated Microbial Genomes & Microbiomes^41^ (IMG/M) database using fairly stringent thresholds of 70% alignment over the query and 90% nucleotide identity. This search yielded 35 hits of the R. UCG13 locus in human microbiome datasets, including senior adults, children, and an infant (12-months of age, Ga0169237_00111) (**Supplemental Table 4**). We also found 12 hits for the *B. intestinalis* XGOs locus, all in human microbiome samples except for a single environmental sample from a fracking water sample from deep shales in Oklahoma, USA (81% coverage, 99% identity) (**Extended Data 10, Supplemental Table 4**). XG and other polysaccharides such as guar gum are used in oil industry processes, and genes for guar gum catabolism have previously been found in oil well associated microbial communities^42^. Since most samples searched were non-gut-derived, this demonstrates that XG-degrading R. UCG13 and XGOs-degrading *B. intestinalis* are largely confined to gut samples and can be present across the human lifetime.

### The mouse microbiome harbors a xanthan utilization locus

To investigate the prevalence of XG-degrading populations beyond the human gut microbiome, we used samples from a previous mouse experiment in which animals fed 5% XG showed increased levels of short chain fatty acids propionate and butyrate, suggesting the ability of members of the mouse microbiome to catabolize and ferment XG^43^. After culturing mouse feces from this experiment on XG media and confirming its ability to depolymerize XG, we used metagenomics to characterize the community structure in two samples (M1741 and M737), revealing the presence of a microbial species related to R. UCG13 (AAI values between the human R. UCG13 and the mouse R. UCG13 were 75.7% and 75.2% for M1741 and M737, respectively) as well as a XG locus with strikingly similar genetic architecture to our previously characterized human XG locus (**Extended Data 10, Supplemental Table 5**). Although several genes are well conserved across both the human and mouse isolates, we observed significant divergence in the sequences of the respective *Ru*GH5a proteins that, based on data with the human locus, initiate XG depolymerization. Specifically, this divergence was more pronounced in the non-catalytic and non-CBM portions of the protein suggesting that while the XG-hydrolyzing functions have been maintained, other domains may be more susceptible to genetic drift. As with the human *Ru*GH5a, recombinant versions of the mouse *Ru*GH5a were able to hydrolyze XG (**Extended Data 10**) but did not show significant activity on a panel of other polysaccharides. These data suggest that the R. UCG13 XG locus is more broadly present in mammalian gastrointestinal microbiomes and can at least be recovered through XG-feeding.

### Prospectus

Our results demonstrate the existence of a multi-phylum food chain in response to XG that appears to have driven the colonization and expansion of R. UCG13 and *B. intestinalis* in industrialized human microbiomes. The absence of these XG-degraders in pre-industrialized microbiomes and their variable presence across post-industrialized populations suggests that XG-driven modulation of human microbiomes may be an ongoing process. The wide range in levels of XG consumption, variable presence of XG-degrading microbes across human populations, and our finding that R. UCG13 can colonize infants at an early age highlight the profound impacts that XG may be having on the assembly, stability, and evolution of industrialized human microbiomes.

The discovery of XG loci in an environmental sample and mouse microbiome, raises ecological questions about the transfer and evolution of XG utilization between host and non-host associated environments. Although the mouse microbiome with a XG locus could have been exposed to XG through herbivory of *X. campestris* infected plants, mice are affiliated with human activities as pests and XG is used as a food additive in various domesticated animal foodstuffs (e.g. in calf milk replacers^44^), further solidifying a link between these loci and human activities. Since XG is a naturally biosynthesized exopolysaccharide, it is also intriguing to speculate about the role of R. UCG13’s XG locus with respect to exopolysaccharides that other microbes may be producing locally in the gut.

While many questions remain about the ecological, functional, and health-relevant impacts of XG on the human microbiome, our study provides strong evidence that food additives should not be considered inert and can be drivers of microbiome ecology with potentially broad impacts.

## Acknowledgments

We gratefully acknowledge Stephanie Theide for growth curve analysis suggestions and Tina Johannessen and Alexsander Lysberg for help with Nanopore metagenomics. We thank the University of Michigan Proteomics Resource Facility, Microbiome Core, and Natural Products Discovery Core for their support in completion of this project. We are grateful for support from the US National Institutes of Health (DK118024, DK125445 to ECM and UL1TR002240 in support of MPO) and the Research Council of Norway (FRIPRO program, PBP and SLLR: 250479, LHH: 302639). We thank the University of Michigan Center for Gastrointestinal Research (UMCGR), (NIDDK 5P30DK034933) for financial support with proteomics. The work conducted by the U.S. Department of Energy Joint Genome Institute, a DOE Office of Science User Facility, is supported under Contract No. DE-AC02-05CH11231.

## Methods

### Isolation, culture, and phylogenetic analysis of xanthan degrading cultures

The original culture was isolated from a survey of 80 healthy adults using a bacterial culture strategy designed to enrich for members of the Gram-negative Bacteroidetes, a phylum that generally harbors numerous polysaccharide-degrading enzymes^23^. The original culture was the only XG-degrading culture isolated from this initial survey, likely due to its bias for Bacteroidetes. For subsequent surveys and further culturing fecal samples were collected into pre-reduced phosphate buffered saline, then transferred to an anaerobic chamber (10% H2, 5% CO2, and 85% N2; Coy Manufacturing, Grass Lake, MI) maintained at 37°C. Fecal suspensions were used to inoculate cultures and passaged using partially Defined Medium (DM), which was generally prepared as a 2x stock then mixed 1:1 with 10 mg/mL carbon source (e.g. xanthan gum). Each L of prepared DM medium (pH=7.2) contained 13.6 g KH2PO4 (Fisher, P284), 0.875 g NaCl (Sigma, S7653), 1.125 g (NH4)2SO4 (Fisher, A702), 2 mg each of adenine, guanine, thymine, cytosine, and uracil (Sigma, A2786, G11950, T0895, C3506, U1128, prepared together as 100x solution), 2 mg of each of the 20 essential amino acids (prepared together as 100x solution), 1 mg vitamin K3 (menadione, Sigma M5625), 0.4 mg FeSO4 (Sigma, 215422), 9.5 mg MgCl2 (Sigma, M8266), 8 mg CaCl2 (Sigma, C1016), 5 µg Vitamin B12 (Sigma, V2876), 1 g L-cysteine, 1.2 mg hematin with 31 mg histidine (prepared together as 1,000x solution), 1 mL of Balch’s vitamins, 1 mL of trace mineral solution, and 2.5 g beef extract (Sigma, B4888).

Each L of Balch’s vitamins was prepared with 5 mg *p*-Aminobenzoic acid, 2 mg folic acid (Sigma, F7876), 2 mg biotin (Sigma, B4501), 5 mg nicotinic acid (Sigma, N4126), 5 mg calcium pantothenate (Sigma, P2250), 5 mg riboflavin (Sigma, R7649), 5 mg thiamine HCl (Sigma, T4625), 10 mg pyridoxine HCl, 0.1 mg cyanocobalamin, 5 mg thioctic acid. Prepared Balch’s vitamins adjusted to pH 7.0, filter sterilized with 0.22 µm PES filters, and stored in the dark at 4 C.

Each L of trace mineral solution was prepared with 0.5 g EDTA (Sigma, ED4SS), 3 g MgSO4*7H2O, 0.5 g MnSO4*H2O, 1 g NaCl (Sigma, S7653), 0.1 g FeSO4*7H2O (Sigma, 215422), 0.1 g CaCl2, 0.1 g ZnSO4*7H2O, 0.01 g CuSO4*5H2O, 0.01 g H3BO3 (Sigma, B6768), 0.01 g Na2MoO4*2H2O, 0.02 g NiCl2*6H2O. Prepared trace mineral solution was adjusted to pH 7.0, filter sterilized with 0.22 µm PES filters, and stored at room temperature. Samples that showed growth on xanthan gum, as evidenced by loss of viscosity and increased culture density, were subcultured 10 times by diluting an active culture 1:100 into fresh DM-XG medium. For the original culture, multiple samples were stored for gDNA extraction and analysis while for the larger sample set, samples were stored after 10 passages; samples were harvested by centrifugation, decanted, and stored at −20 C until further processing).

Frozen cell pellets were resuspended in 500 µL Buffer A (200 mM NaCl, 200 mM Tris-HCl, 20 mM EDTA) and combined with 210 µL SDS (20% w/v, filter-sterilized), 500 µL phenol:chloroform (alkaline pH), and ∼250 µL acid-washed glass beads (212-300 µm; Sigma).

Samples were bead beaten on high for 2-3 minutes with a Mini-BeadBeater-16 (Biospec Products, USA), then centrifuged at 18,000 *g* for 5 mins. The aqueous phase was recovered and mixed by inversion with 500 µL of phenol:chloroform, centrifuged at 18,000 *g* for 3 mins, and the aqueous phase was recovered again. The sample was mixed with 500 µL chloroform, centrifuged, and then the aqueous phase was recovered and mixed with 0.1 volumes of 3 M sodium acetate (pH 5.2) and 1 volume isopropanol. The sample was stored at −80 C for ≥30 mins, then centrifuged at ≥20,000 *g* for 20 mins at 4 C. The pellet was washes with 1 mL room temperature 70% ethanol, centrifuged for 3 mins, decanted, and allowed to air dry before resuspension in 100 µL sterile water. Resulting samples were additionally purified using the DNeasy Blood & Tissue Kit (QIAGEN, USA).

Illumina sequencing, including PCR and library preparation, were performed by the University of Michigan Microbiome Core as described by Kozich et al^45^. Barcoded dual-index primers specific to the 16S rRNA V4 region were used to amplify the DNA. PCR reactions consisted of 5 µL of 4 µM equimolar primer set, 0.15 µL of AccuPrime Taq DNA High Fidelity Polymerase, 2 µL of 10x AccuPrime PCR Buffer II (Thermo Fisher Scientific, catalog no. 12346094), 11.85 µL of PCR-grade water, and 1 µL of DNA template. The PCR conditions used consisted of 2 min at 95°C, followed by 30 cycles of 95°C for 20 s, 55°C for 15 s, and 72°C for 5 min, followed by 72°C for 10 min. Each reaction was normalized using the SequalPrep Normalization Plate Kit (Thermo Fisher Scientific, catalog no. A1051001), then pooled and quantified using the Kapa Biosystems Library qPCR MasterMix (ROX Low) Quantification kit for Illumina platforms (catalog no. KK4873). After confirming the size of the amplicon library using an Agilent Bioanalyzer and a high-sensitive DNA analysis kit (catlog no. 5067-4626), the amplicon library was sequenced on an Ilumina MiSeq platform using the 500 cycle MiSeq V2 Reagent kit (catalog no. MS-102-2003) according to the manufacturer’s instructions with with modifications of the primer set with custom read 1/read 2 and index primers added to the reagent cartridge. The “Preparing Libraries for Sequencing on the MiSeq” (part 15039740, Rev. D) protocol was used to prepare libraries with a final load concentration of 5.5 pM, spiked with 15% PhiX to create diversity within the run.

Sequencing FASTQ files were analyzed using mothur (v.1.40.5)^46^ using the Silva reference database^11^. OTUs with the same genus were combined and displayed using R^47^ with the packages reshape2^48^, RColorBrewer^49^, and ggplot2^50^.

### Dilution to extinction experiment

An overnight culture was serially diluted in 2x DM. Serial dilutions were split into two 50 mL tubes and mixed 1:1 with either 10 mg/mL xanthan gum or 10 mg/mL monosaccharide mixture (4 mg/mL glucose, 4 mg/mL mannose, 2 mg/mL sodium glucuronate), both of which also had 1 mg/mL L-cysteine. Each dilution and carbon source was aliquoted to fill a full 96-well culture plate (Costar 3370) with 200 µL per well. Plates were sealed with Breathe-Easy gas permeable sealing membrane for microtiter plates (Diversified Biotech, cat #BEM-1). Microbial growth was measured at least 60 hours by monitoring OD_600_ using a Synergy HT plate reader (Biotek Instruments) and BIOSTACK2WR plate handler (Biotek Instruments)^51^.

Maximum OD for each substrate was measured for each culture. Full growth on substrates was conservatively defined as a maximum OD_600_ of >0.7. For each unique 96 well plate of substrate and dilution factor, the fraction of wells exhibiting full growth was calculated. Fractional growth was plotted against dilution factor for each substrate. Data were fit to the Hill equation by minimizing squared differences between the model and experimental values using Solver (GRG nonlinear) in Excel. For each experiment, a 50% growth dilution factor (GDF 50) was calculated for each substrate at which half of the wells would be predicted to exhibit full growth.

### Metagenomic analysis

Seven samples (15-mL) were collected at four time points (Extended Data 3; referred to as T1, T2, T3 and T4) during growth of two biological replicates of the original XG-degrading culture. Cells were harvested by centrifugation at 14,000 × g for 5 min and stored a −20 °C until further use. A phenol:chloroform:isoamyl alcohol and chloroform extraction method was used to obtain high molecular weight DNA as previously described^52^. The gDNA was quantified using a Qubit™ fluorimeter and the Quant-iT™ dsDNA BR Assay Kit (Invitrogen, USA), and the quality was assessed with a NanoDrop One instrument (Thermo Fisher Scientific, USA). Samples were subjected to metagenomic shotgun sequencing using the Illumina HiSeq 3000 platform at the Norwegian Sequencing Center (NSC, Oslo, Norway). Samples were prepared with the TrueSeq DNA PCR-free preparation and sequenced with paired ends (2 × 150 bp) on one lane. Quality trimming of the raw reads was performed using Cutadapt^53^ v1.3, to remove all bases on the 3′-end with a Phred score lower than 20 and exclude all reads shorter than 100 nucleotides, followed by a quality filtering using the FASTX-Toolkit v.0.0.14 (http://hannonlab.cshl.edu/fastx_toolkit/). Retained reads had a minimum Phred score of 30 over 90% of the read length. Reads were co-assembled using metaSPAdes^54^ v3.10.1 with default parameters and k-mer sizes of 21, 33, 55, 77 and 99. The resulting contigs were binned with MetaBAT^55^ v0.26.3 in “very sensitive mode”. The quality (completeness, contamination, and strain heterogeneity) of the metagenome assembled genomes (MAGs) was assessed by CheckM^56^ v1.0.7 with default parameters. Contigs were submitted to the Integrated Microbial Genomes and Microbiomes system for open reading frames (ORFs) prediction and annotation^57^. Additionally, the resulting ORF were annotated for CAZymes using the CAZy annotation pipeline^58^. This MAG collection was used as a reference database for mapping of the metatranscriptome data, as described below. Taxonomic classifications of MAGs were determined using both MiGA^59^ and GTDB-Tk^60^.

Human fecal samples (20) from a second enrichment experiment (unbiased towards the cultivation of Bacteroides) as well as two enrichments with mouse fecal samples were processed for gDNA extraction and library preparation exactly as described above. Metagenomic shotgun sequencing was conducted on two lanes of both Illumina HiSeq 4000 and Illumina HiSeq X Ten platforms (Illumina, Inc.) at the NSC (Oslo, Norway), and reads were quality trimmed, assembled and binned as described above. Open reading frames were annotated using PROKKA^61^ v1.14.0 and resulting ORFs were further annotated for CAZymes using the CAZy annotation pipeline and expert human curation^58^. Completeness, contamination, and taxonomic classifications for each MAG were determined as described above. AAI comparison between the human R. UCG13 and the R. UCG13 found in the two mouse samples was determined using CompareM (https://github.com/dparks1134/CompareM).

Extracted DNA from a second enrichment experiment on XG using the original culture was prepared for long-reads sequencing using Oxford Nanopore Technologies (ONT) Ligation Sequencing Kit (SQK-LSK109) according to the manufacture protocol. The DNA library was sequenced with the ONT MinION Sequencer using a R9.4 flow cell. The sequencer was controlled by the MinKNOW software v3.6.5 running for 6 hours on a laptop (Lenovo ThinkPad P73 Xeon with data stored to 2Tb SSD), followed by base calling using Guppy v3.2.10 in ‘fast’ mode. This generated in total 3.59 Gb of data. The Nanopore reads were further processed using Filtlong v0.2.0 (https://github.com/rrwick/Filtlong), discarding the poorest 5% of the read bases, and reads shorter than 1000 bp.

The quality processed Nanopore long-reads were assembled using CANU^62^ v1.9 with the parameters *corOutCoverage=10000 corMinCoverage=0 corMhapSensitivity=high genomeSize=5m redMemory=32 oeaMemory=32 batMemory=200*. An initial polishing of the generated contigs were carried out using error-corrected reads from the assembly with minimap2^63^ v2.17 *-x map-ont* and Racon^64^ v1.4.14 with the argument *--include-unpolished*. The racon-polished contigs were further polished using Medaka v1.1.3 (https://github.com/nanoporetech/medaka), with the commands *medaka_consensus --model r941_min_fast_g303_model.hdf5*. Finally, Minimap2 *-ax sr* was used to map quality processed Illumina reads to the medaka-polished contigs, followed by a final round of error correction using Racon with the argument *--include-unpolished*. Circular contigs were identified by linking the contig identifiers in the polished assembly back to *suggestCircular=yes* in the initial contig header provided by CANU. These contigs were quality checked using CheckM^56^ v1.1.3 and BUSCO^65^ v4.1.4. Circular contigs likely to represent chromosomes (> 1 Mbp) were further gene-called and functionally annotated using PROKKA^61^ v1.13 and taxonomically classified using GTDB-tk^60^ v1.4.0 with the *classify_wf* command. Barrnap v0.9 (https://github.com/tseemann/barrnap) was used to predict ribosomal RNA genes. Average nucleotide Identity (ANI) was measured between the short-reads and long-reads MAGs using FastANI^66^ v1.1 with default parameters. Short-reads MAGs were used as query while long-reads MAGs were set as reference genomes. Short-reads MAG1 showed an Average Nucleotide Identity (ANI) of 99.98% with the long-reads ONT_Circ01, while short-reads MAG2 showed an ANI of 99.99% with the long-reads ONT_Circ02 (**Supplemental Table 2**). Phylogenetic analysis revealed that ONT_Circ02 encoded four complete 16S rRNA operons, three of which were identical to the aforementioned R. UCG13 OTU.

### Temporal metatranscriptomic analysis of the original XG-degrading community

Cell pellets from 6 mL samples collected at T1-T4 during growth of two biological replicates of the original XG-degrading culture were supplemented with RNAprotect Bacteria Reagent (Qiagen, USA) following the manufacturer’s instructions and kept at −80 °C until RNA extraction. mRNA extraction and purification were conducted as described in Kunath et al.^67^. Samples were processed with the TruSeq stranded RNA sample preparation, which included the production of a cDNA library, and sequenced on one lane of the Illumina HiSeq 3000 system (NSC, Oslo, Norway) to generate 2 × 150 paired-end reads. Prior to assembly, RNA reads were quality filtered with Trimmomatic^68^ v0.36, whereby the minimum read length was required to be 100 bases and an average Phred threshold of 20 over a 10 nt window, and rRNA and tRNA were removed using SortMeRNA^69^ v.2.1b. Reads were pseudo-aligned against the metagenomic dataset using kallisto pseudo –pseudobam^70^. Of the 58089 ORFs (that encode proteins with > 60 aa) identified from the metagenome of the original XG-degrading community, 7549 (13%) were not found to be expressed, whereas 50540 (87%) were expressed, resulting in a reliable quantification of the expression due to unique hits (reads mapping unambiguously against one unique ORF).

### Neutral Monosaccharide analysis

The hot-phenol extraction method originally described by Massie & Zimm^71^ and modified by Nie^72^ was used for collecting and purifying the polysaccharides remaining at different timepoints. Samples were heated to 65 °C for 5 mins, combined with an equal volume of phenol, incubated at 65 °C for 10 mins, then cooled to 4 °C and centrifuged at 4 °C for 15 min at 12,000 *g*. The upper aqueous layer was collected and re-extracted using the same procedure, dialyzed extensively against deionized water (2000 Da cutoff), and freeze-dried. Neutral monosaccharide composition was obtained using the method described by Tuncil et al.^73^ Briefly, sugar alditol acetates were quantified by gas chromatography using a capillary column SP-2330 (SUPELCO, Bellefonte, PA) with the following conditions: injector volume, 2 μl; injector temperature, 240 °C; detector temperature, 300 °C; carrier gas (helium), velocity 1.9 meter/second; split ratio, 1:2; temperature program was 160 °C for 6 min, then 4 °C/min to 220 °C for 4 min, then 3 °C/min to 240 °C for 5 min, and then 11 °C/min to 255 °C for 5 min.

### Thin Layer Chromatography for Localization of Enzyme Activity

Overnight cultures were harvested at 13,000 *g* for 10 minutes. Supernatant fractions were prepared by vacuum filtration through 0.22 µm PES filters. Cell pellet fractions were prepared by decanting supernatant, washing with phosphate buffered saline (PBS), spinning at 13,000 *g* for 3 mins, decanting, and resuspending in PBS. Intracellular fractions were prepared by taking cell pellet fractions and bead beating for 90 s with acid-washed glass beads (G1277, Sigma) in a Biospec Mini Beadbeater. Lysed culture fractions were prepared by directly bead beating unprocessed culture.

Each culture fraction was mixed 1:1 with 5 mg/mL xanthan gum and incubated at 37 C for 24 hours. Negative controls were prepared by heating culture fractions to 95 C for 15 mins, then centrifuging at 13,000 *g* for 10 mins before the addition of xanthan gum. All reactions were halted by heating to ≥85 C for 15 mins, then spun at 20,000 *g* for 15 mins at 4 C. Supernatants were stored at −20 C until analysis by thin layer chromatography.

3 µL sample were spotted twice onto a 10×20 cm thin layer chromatography plate (Millipore TLC Silica gel 60, 20×20cm aluminum sheets), with intermediate drying using a Conair 1875 hairdryer. Standards included malto-oligosaccharides of varying lengths (Even: 2, 4, 6, Odd: 1, 3, 5, 7), glucuronic acid, and mannose. Standards were prepared at 10 mM and 3 uL of each was spotted onto the TLC plate. Plates were run in ∼100 mL of 2:1:1 butanol, acetic acid, water, dried, then run an additional time. After drying, plates were incubated in developing solution (100 mL ethyl acetate, 2 g diphenylamine, 2 mL aniline, 10 mL of ∼80% phosphoric acid, 1 mL of ∼38% hydrochloric acid) for ∼30 seconds, then dried, and developed by holding over a flame until colors were observed.

### Proteomic analysis

Approximately 1 L of xanthan gum culture was grown until it had completely liquified (∼2-3 days). Supernatant was collected by centrifuging at 18,000 *g* and vacuum filtering through a 0.2 µm PES filter. 4M ammonium sulfate was added to 200-400 mL of filtrate to a final concentration of 2.4M and incubated for 30-60 mins at RT or, for one sample, overnight at 4 C. Precipitated proteins were harvested by centrifugation at 18,000 *g* for 30-60 mins, then resuspended in 50 mM sodium phosphate (pH 7.5). Three different fractionation protocols were followed, but after every fractionation step, active fractions were identified by mixing ∼500 µL with 10 mg/mL xanthan and incubating at 37 C overnight; active-fractions were identified by loss of viscosity or production of xanthan oligosaccharides as visualized by TLC (method previously described).

1. Resuspended protein was filtered and applied to a HiTrapQ column, running a gradient from 0-100% B (Buffer A: 50 mM sodium phosphate, pH 7.5; Buffer B: 50 mM sodium phosphate, 1 M NaCl, pH 7.5). Active fractions were pooled and concentrated with a 10 kDa MWCO centricon and injected onto an S-200 16/60 column equilibrated in 50 mM sodium phosphate, 200 mM NaCl, pH 7.5. The earliest fractions to elute with significant A280 absorbance were also the most active fractions; these were pooled and submitted for proteomics.
2. Resuspended protein was filtered and applied to an S-500 column equilibrated in 50 mM sodium phosphate, 200 mM NaCl, pH 7.5. Active fractions eluted in the middle of the separation were pooled and submitted for proteomics.
3. Resuspended protein was filtered and applied to an S-500 column equilibrated in 50 mM sodium phosphate, 200 mM NaCl, pH 7.5. Pooled fractions were applied to a 20 mL strong anion exchange column running a gradient from 0-100% B (Buffer A: 50 mM sodium phosphate, pH 7.5; Buffer B: 50 mM sodium phosphate, 1 M NaCl, pH 7.5). Active fractions were pooled and applied to a 1 mL weak anion exchange column (ANX) running a gradient from 0-100% B (Buffer A: 50 mM sodium phosphate, 10% glycerol, pH 7.5; Buffer B: 50 mM sodium phosphate, 1 M NaCl, 10% glycerol, pH 7.5). Active fractions were pooled and submitted for proteomics.

Cysteines were reduced by adding 50 ml of 10 mM DTT and incubating at 45 °C for 30 min. Samples were cooled to room temperature and alkylation of cysteines was achieved by incubating with 65 mM 2-Chloroacetamide, under darkness, for 30 min at room temperature. An overnight digestion with 1 ug sequencing grade, modified trypsin was carried out at 37 C with constant shaking in a Thermomixer. Digestion was stopped by acidification and peptides were desalted using SepPak C18 cartridges using manufacturer’s protocol (Waters). Samples were completely dried using vacufuge. Resulting peptides were dissolved in 8 ml of 0.1% formic acid/2% acetonitrile solution and 2 µls of the peptide solution were resolved on a nano-capillary reverse phase column (Acclaim PepMap C18, 2 micron, 50 cm, ThermoScientific) using a 0.1% formic acid/2% acetonitrile (Buffer A) and 0.1% formic acid/95% acetonitrile (Buffer B) gradient at 300 nl/min over a period of 180 min (2-25% buffer B in 110 min, 25-40% in 20 min, 40-90% in 5 min followed by holding at 90% buffer B for 10 min and requilibration with Buffer A for 30 min). Eluent was directly introduced into *Q exactive HF* mass spectrometer (Thermo Scientific, San Jose CA) using an EasySpray source. MS1 scans were acquired at 60K resolution (AGC target=3×10^6^; max IT=50 ms). Data-dependent collision induced dissociation MS/MS spectra were acquired using Top speed method (3 seconds) following each MS1 scan (NCE ∼28%; 15K resolution; AGC target 1×10^5^; max IT 45 ms).

Proteins were identified by searching the MS/MS data against a database of all proteins identified in the original culture metagenomes using Proteome Discoverer (v2.1, Thermo Scientific). Search parameters included MS1 mass tolerance of 10 ppm and fragment tolerance of 0.2 Da; two missed cleavages were allowed; carbamidimethylation of cysteine was considered fixed modification and oxidation of methionine, deamidation of asparagine and glutamine were considered as potential modifications. False discovery rate (FDR) was determined using Percolator and proteins/peptides with an FDR of ≤1% were retained for further analysis.

### Plasmid Design and Protein Purification

Plasmid constructs to produce recombinant proteins were made with a combination of synthesized DNA fragments (GenScript Biotech, Netherlands) and PCR amplicons using extracted culture gDNA as a template. In general, sequences were designed to remove N-terminal signaling peptides and to add a histidine tag for immobilized metal affinity chromatography (IMAC) (in many cases using the Lucigen MA101-Expresso-T7-Cloning-&-Expression-System). Plasmid assembly and protein sequences are described in **Supplemental Table 6**.

Constructs were transformed into HI-Control BL21(DE3) cells and single colonies were inoculated in 5 mL overnight LB cultures at 37°C. 5 mL cultures were used to inoculate 1 L of Terrific Broth (TB) with selective antibiotic, grown to OD ∼0.8-1.1 at 37°C, and induced with 250 µM IPTG. *B. intestinalis* enzymes were expressed at RT, while R. UCG13 enzymes were expressed at 18°C overnight. Cells were harvested by centrifugation and pellets were stored at −80°C until further processing. Proteins were purified using standard IMAC purification procedures employing sonication to lyse cells. R. UCG13 proteins were purified using 50 mM sodium phosphate and 300 mM sodium chloride at pH 7.5; *B. intestinalis* proteins were purified using 50 mM Tris and 300 mM sodium chloride at pH 8.0. All proteins were eluted from cobalt resin using buffer with the addition of 100 mM imidazole, then buffer exchanged to remove imidazole using Zeba 2 mL 7kDa MWCO desalting columns. Protein concentrations were determined by measuring A280 and converting to molarity using calculated extinction coefficients.

### Characterization and isolation of Xanthan gum degradation products

In general, pentameric xanthan oligosaccharides were produced by incubating ≥0.1 mg/mL *Ru*GH5a with 5 mg/mL xanthan gum in PBS in approximately 1L total volume. For xanthan tetrasaccharides, ∼0.5 U/mL of Xanthan lyase (E-XANLB, Megazyme) was included. After incubating 2-3 days at 37 °C to allow complete liquefication, reactions were heat-inactivated, centrifuged at ≥10,000 *g* for 30 mins, and the supernatant was vacuum filtered through 0.22 µm PES sterile filters. Supernatants were loaded onto a column containing ∼10 g of graphitized carbon (Supelclean™ ENVI-Carb™, 57210-U Supelco), washed extensively with water to remove salt and unbound material, then eluted in a stepwise fashion with increasing concentrations of acetonitrile. Fractions were dried, weighed, and analyzed by LC-MS and fractions that contained the most significant yield of desired products were combined. Highly pure products were obtained by reconstituting samples in 50% water:acetonitrile and applying to a Luna® 5 µm HILIC 200 Å LC column (250 x 10 mm) (00G-4450-N0, Phenomenex). A gradient was run from 90-20% acetonitrile, with peaks determined through a combination of evaporative light scattering, UV, and post-run analytical LC-MS (Agilent qToF 6545) of resulting fractions.

NMR spectra were collected using an Agilent 600 NMR spectrometer (^1^H: 600 MHz, ^13^C: 150 MHz) equipped with a 5 mm DB AUTOX PFG broadband probe and a Varian NMR System console. All data analysis was performed using MestReNova NMR software. All chemical shifts were referenced to residual solvent peaks [^1^H (D_2_O): 4.79 ppm].

### Enzyme Reaction Analysis

All enzyme reactions were carried out in 15-25 mM sodium phosphate buffer, 100-150 mM sodium chloride, and sometimes included up to 0.01 mg/mL bovine serum albumin (B9000S, NEB) to limit enzyme adsorption to pipettes and tubes. All R. UCG13 or *B. intestinalis* enzymes were tested at concentrations from 1-10 µM. Cellobiose reactions were tested using 1 mM cellobiose at pH 7.5, while all other reactions used 2.5 mg/mL pentasaccharide (produced using *Ru*GH5a) and were carried out at pH 6.0. Reactions were incubated overnight at 37°C, halted by heating at ≥ 95°C for 5-10 minutes, and centrifugation at ≥20,000 *g* for 10 mins. Supernatants were mixed with 4 parts acetonitrile to yield an 80% acetonitrile solution, centrifuged for 10 mins at ≥20,000 *g*, and transferred into sample vials. 15 µL of each sample was injected onto a Luna® Omega 3 µm HILIC 200 Å LC column (100 x 4.6 mm) (00D-4449-E0, Phenomenex). An Agilent 1290 Infinity II HPLC system was used to separate the sample using solvent A (100% water, 0.1% formic acid) and solvent B (95% acetonitrile, 5% water, with 0.1% formic acid added) at a flow rate of 0.4 mL/min. Prior to injection and following each sample the column was equilibrated with 80% B. After injection, samples were eluted with a 30 minute isocratic step at 80% B, a 10 minute gradient decreasing B from 80% to 10%, and a final column wash for 2 min at 10% B. Spectra were collected in negative mode using an Agilent 6545 LC/Q-TOF.

### Kinetics of RuGH5a (BCA Assay)

Lyase-treated xanthan gum was generated by mixing 5 mg/mL xanthan gum with 0.5 U/mL of *Bacillus* sp. Xanthan lyase (E-XANLB, Megazyme) in 30 mM potassium phosphate buffer (pH 6.5). After incubating overnight at 37 °C, an additional 0.5 U/mL of xanthan lyase was added. Both lyase-treated and native xanthan gum were dialyzed extensively against deionized water, heated in an 80 °C water bath to inactivate the lyase, and centrifuged at 10,000 *g* for 20 mins to remove particulate. Supernatants were collected and stored at 4 °C until use.

Kinetic measurements were conducted using a slightly modified version of the low-volume bicinchoninic acid (BCA) assay for glycoside hydrolases used by Arnal et al^74^. Briefly, AEX and SEC purified *Ru*GH5a was diluted to a 10x stock of 5 µM enzyme, 50 mM sodium phosphate, 300 mM sodium chloride, and 0.1 mg/mL bovine serum albumin, pH7.5. Reactions were 20 µL of enzyme stock mixed with 180 µL of various concentrations of xanthan gum. Negative controls were conducted with heat-inactivated enzyme stock. Timepoints were taken by quenching reactions with dilute, ice-cold, BCA working reagent. Reactions and controls were run with 4 independent replicates and compared to a glucose standard curve. Enzyme released reducing sugar was calculated by subtracting controls from reaction measurements.

### Growth curves of isolates on XG oligos

Pure isolates from the xanthan culture were obtained by streaking an active culture onto a variety of agar plates including LB and brain heart infusion with the optional addition of 10% defibrinated horse blood (Colorado Serum Co.) and gentamicin (200 µg/mL). After passaging isolates twice on agar plates, individual colonies were picked and grown overnight in tryptone-yeast extract-glucose (TYG) broth medium, then stocked by mixing with 0.5 volumes each of TYG and molecular biology grade glycerol and storing at −80 °C.

DM without beef extract (DM^-BE^), with the addition of a defined carbon source, was used to test isolates for growth on xanthan oligosaccharides. Some isolates (e.g. *Parabacteroides distasonis*) required the inclusion of 5 mg/mL beef extract (Sigma, B4888) to achieve robust growth on simple monosaccharides; in these cases, beef extract was included across all carbon conditions. Unless otherwise specified, carbon sources were provided at a final concentration of 5 mg/mL. Isolates were grown overnight in TYG media, subcultured 1:50 into DM^-BE^-glucose and grown overnight, then subcultured 1:50 into DM^-BE^ with either various carbon sources. Final cultures were monitored for growth by measuring increase in absorbance (600 nm) using 96-well plates as previously described.

### qPCR and RNA-seq on B. intestinalis and original community

For qPCR, *B. intestinalis* was grown as before but cells were harvested by centrifugation at mid-exponential phase, mixed with RNA Protect (QIAGEN), and stored at −80 °C until further processing. At collection, average OD_600_ values were ∼0.8 and ∼0.6 for glucose- and oligosaccharide-grown cultures, respectively. RNeasy mini kit buffers (QIAGEN) were used to extract total RNA, purified with RNA-binding spin columns (Epoch), treated with DNase I (NEB), and additionally purified using the RNeasy mini kit. SuperScript III reverse transcriptase and random primers (Invitrogen) were used to perform reverse transcription. Target transcript abundance in the resulting cDNA was quantified using a homemade qPCR mix as described previously^75^ and gene-specific primers (**Supplemental Table 7**). Each 20 uL reaction contained 1X Thermopol Reaction Buffer (NEB), 125uM dNTPs, 2.5mM MgSO4, 1X SYBR Green I (Lonza), 500nM gene specific or 65nM 16S rRNA primer and 0.5 units Hot Start Taq Polymerase (NEB), and 10ng of template cDNA. Results were processed using the ddCT method in which raw values were normalized to 16S rRNA values, then xanthan oligosaccharide values were compared to those from glucose to calculate fold-change in expression.

For RNA-seq, total RNA was used from the *B. intestinalis* growths used for qPCR. For the community grown on XG or PGA, 5 mL cultures of DM-XG or DM-PGA were inoculated with a 1:100 dilution of a fully liquified DM-XG culture. PGA cultures were harvested at mid-log phase at OD_600_ ∼0.85 whereas XG cultures were harvested at late-log phase at OD600 ∼1.2 to allow liquification of XG, which was necessary to extract RNA from these cultures. As before, cultures were harvested by centrifugation, mixed with RNA Protect (Qiagen) and stored at −80 °C until further processing. RNA was purified as before except that multiple replicates of DM-XG RNA were pooled together and concentrated with Zymo RNA Clean and Concentrator^TM^-25 to reach acceptable concentrations for RNA depletion input. rRNA was depleted twice from the purified total RNA using the MICROB*Express*^TM^ Kit, each followed by a concentration step using the Zymo RNA Clean and Concentrator^TM^-25. About 90% rRNA depletion was achieved for all samples. *B. intestinalis* RNA was sequenced using NovaSeq and community RNA was sequenced using MiSeq. The resulting sequence data was analyzed for differentially expressed genes following a previously published protocol^76^. Briefly, reads were filtered for quality using Trimmomatic v0.39^68^. Reads were aligned to the each genome using BowTie2 v2.3.5.1^77^. For the *Bacteroides intestinalis* transcriptome reads were aligned to its genome, while for the community data reads were aligned to either the *B. intestinalis* genome or the closed Ruminococcaceae UCG-13 metagenome assembled genome (MAG). Reads mapping to gene features were counted using htseq-count (release_0.11.1)^78^. Differential expression analysis was performed using the edgeR v3.34.0 package in R v.4.0.2 (with the aid of Rstudio v1.3.1093). The TMM method was used for library normalization^79^. Coverage data was visualized using Integrated Genome Viewer (IGV)^80^.

### Extended metagenome analysis/comparison methodology

Individual MAGs in each sample were searched by BlastP for the presence of proteins similar to those encoded by the XG-degrading PUL of R. UCG13 and *B. intestinalis*. This was done using the amino acid sequences of the proteins in the R. UCG13 and *B. intestinalis* PULs as the search homologs; both BlastP probes were searched against all the individual MAGs in the different samples with the default threshold e-value of 1e-5.

### Looking for R. UCG13 and B. intestinalis. XG Loci in Metagenomes

Available cohorts of human gut metagenomic sequence data (National Center for Biotechnology Information projects: PRJNA422434^81^, PRJEB10878^82^, PRJEB12123^83^, PRJEB12124^84^, PRJEB15371^85^, PRJEB6997^86^, PRJDB3601^87^, PRJNA48479^88^, PRJEB4336^89^, PRJEB2054^90^, PRJNA392180^91^, and PRJNA527208^92^) were searched for the presence of xanthan locus nucleotide sequences from R. UCG13 (92.7 kb) and *B. intestinalis* (17.9kb) using the following workflow: Each xanthan locus nucleotide sequence was used separately as a template and then magic-blast v1.5.0^93^ was used to recruit raw Illumina reads from the available metagenomic datasets with an identity cutoff of 97%. Next, the alignment files were used to generate a coverage map using bedtools v2.29.0^94^ to calculate the percentage coverage of each sample against each individual reference. We considered a metagenomic data sample to be positive for a particular xanthan locus if it had at least 70% of the corresponding xanthan locus nucleotide sequence covered.

The R. UCG13 locus and *B. intestinalis* XG locus were used as the query in a large-scale search against the assembled scaffolds of isolates, metagenome assembled genomes (bins), and metagenomes included into the Integrated Microbial Genomes & Microbiomes (IMG/M) comparative analysis system^41^. Within the LAST software package, version 1066, the ‘lastal’ tool was used with default thresholds to search the 2 loci against 72,491 public high-quality isolate genomes, and 102,860 bins from 13,415 public metagenomes, and 21,762 public metagenomes in IMG/M. Metagenome bins were generated using the binning analysis method described in A. Clum et al^95^.

## Data availability

All sequencing reads have been deposited at the European Nucleotide Archive under BioProject PRJEB44146. All annotated MAGs are publicly available via Figshare (DOIs: 10.6084/m9.figshare.14494602, 10.6084/m9.figshare.14494536, 10.6084/m9.figshare.14494677, 10.6084/m9.figshare.14494683 and 10.6084/m9.figshare.14494689).

---

**Extended Data 1.**
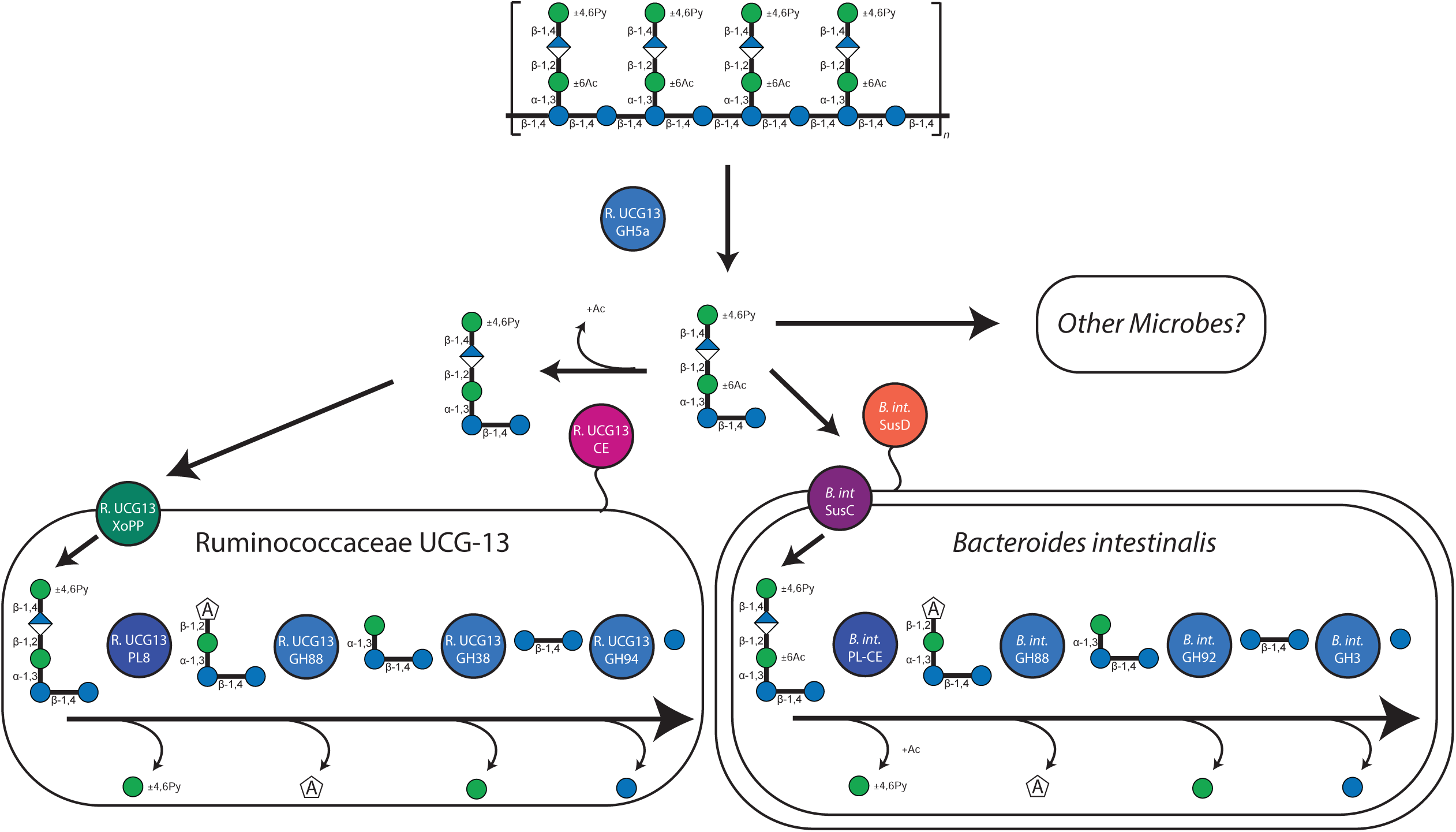
Cellular model of xanthan degradation.

**Extended Data 2.**
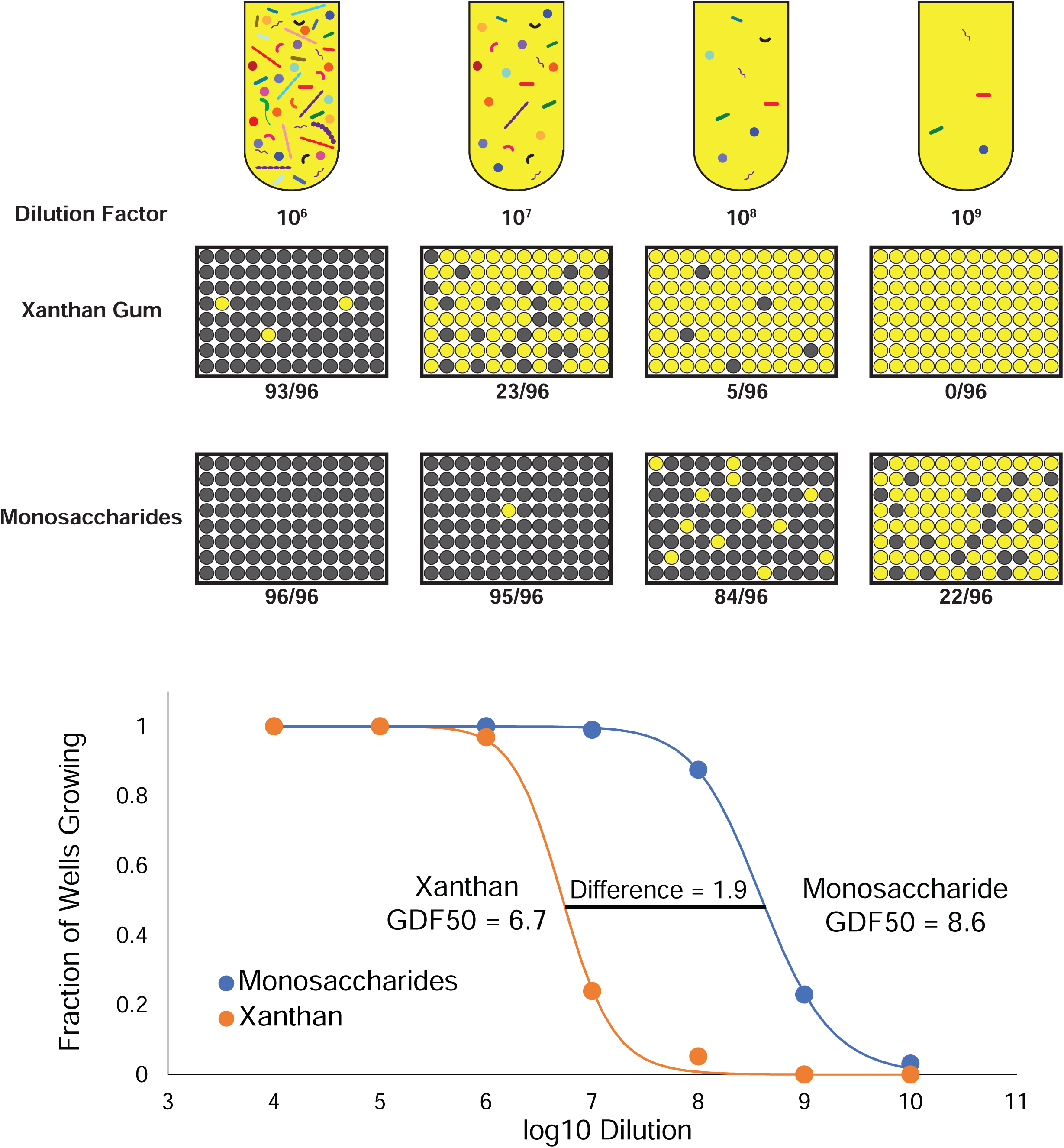
Xanthan degradation is a multi-species phenotype. An active xanthan culture was diluted in 2x defined media without a major carbon source, then divided and diluted 1:1 with either 2x xanthan gum or 2x monosaccharide mix (2:2:1 mannose;glucose;glucuronic acid), then aliquoted into 200 uL cultures in 96-well plates. Each datapoint represents the fraction of cultures (out of 96) growing above OD600 0.7 at each dilution, grown in either xanthan gum or monosaccharide mix media. Data were fit to the Hill equation to calculate a 50% growth dilution factor (GDF 50) at which half of the cultures would grow above OD 0.7. Across 5 independent experiments, there was a GDF50 difference of 1.8 (standard deviation = 0.4, SEM = 0.2). This demonstrates that at comparable dilutions, microbes were present that could grow on monosaccharides but were unable to grow using XG, suggesting that several microbes are required in this media to allow growth on XG.

**Extended Data 3.**
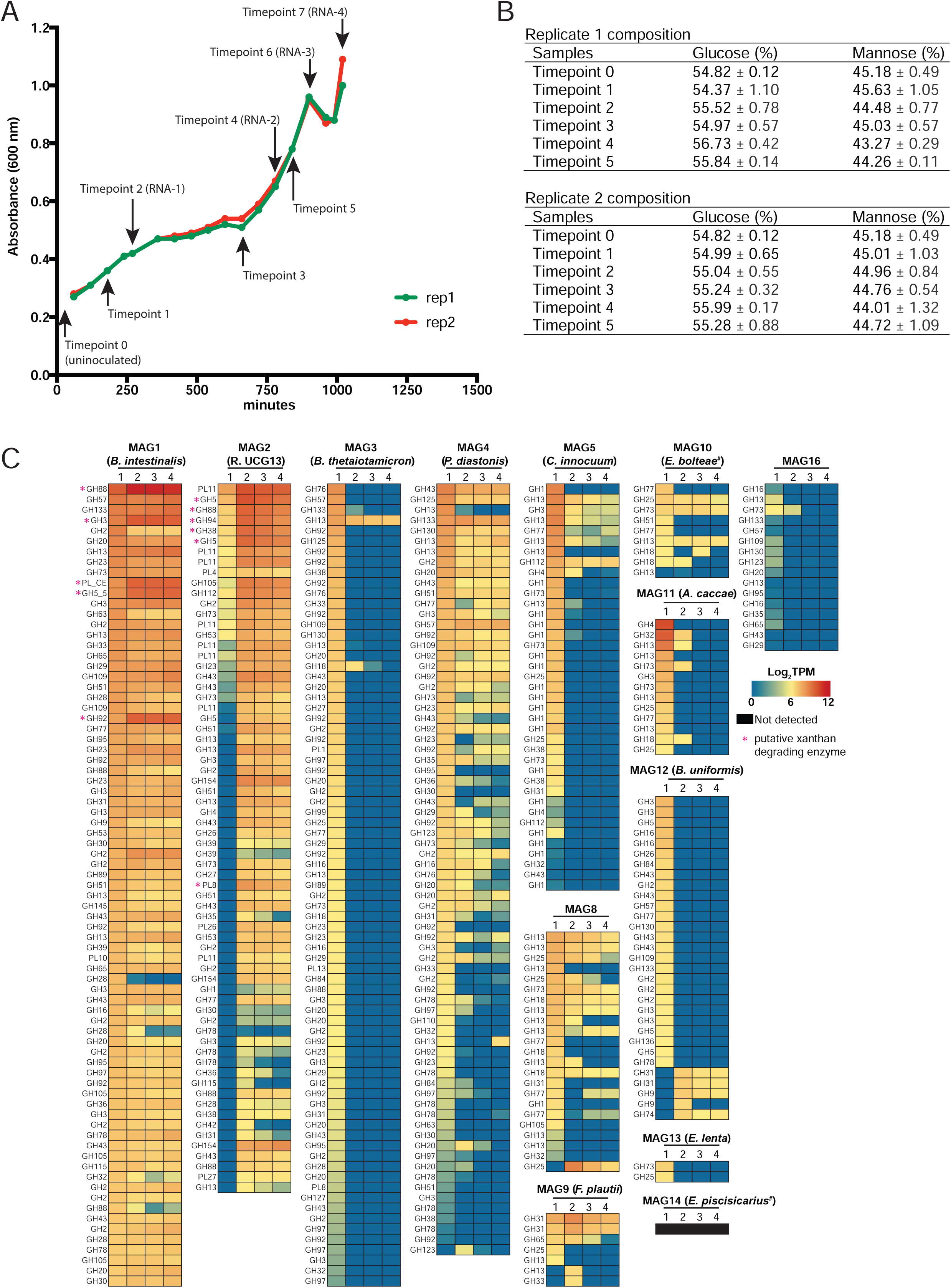
Neutral monosaccharide and metatranscriptomic analysis. **a**, Two replicates of the original xanthan culture were grown and sampled at multiple timepoints for **b**, neutral monosaccharide analysis of residual xanthan gum (n=3, standard deviation shown) and **c**, metatranscriptomic analysis of annotated CAZymes in each of the MAGs (completeness value > 75%) reconstructed from metagenomic data from the enrichment culture. MAG taxonomy (**Supplementary Table 2**) is indicated in parentheticals. An “#” indicates a low AAI%.

**Extended Data 4.**
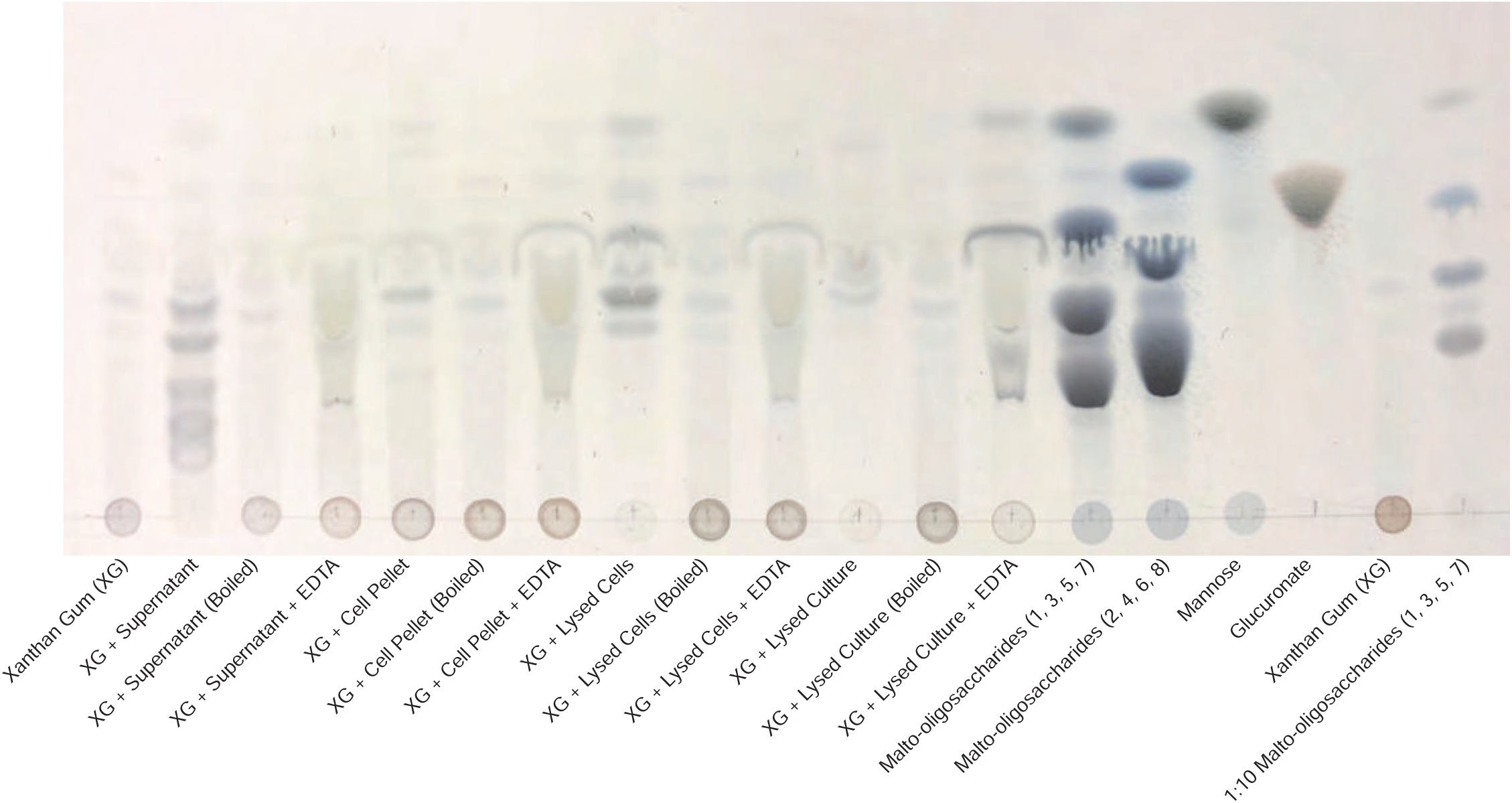
Culture supernatant contains enzymes capable of depolymerizing xanthan gum, while intracellular contents are required for complete saccharification. Thin layer chromatography of xanthan gum incubated with different fractions of an active xanthan gum culture (supernatant, washed cell pellet, lysed cell pellet, or lysed culture). Negative controls were prepared by heating fractions at 95 C for 15 minutes prior to initiating with xanthan gum. EDTA was added to a final concentration of ∼50 mM to determine the necessity of divalent cations for enzyme activity. Strong color development in circles at baseline is undigested polysaccharide while bands that migrated with solvent are digested oligosaccharides and monosaccharides.

**Extended Data 5.**
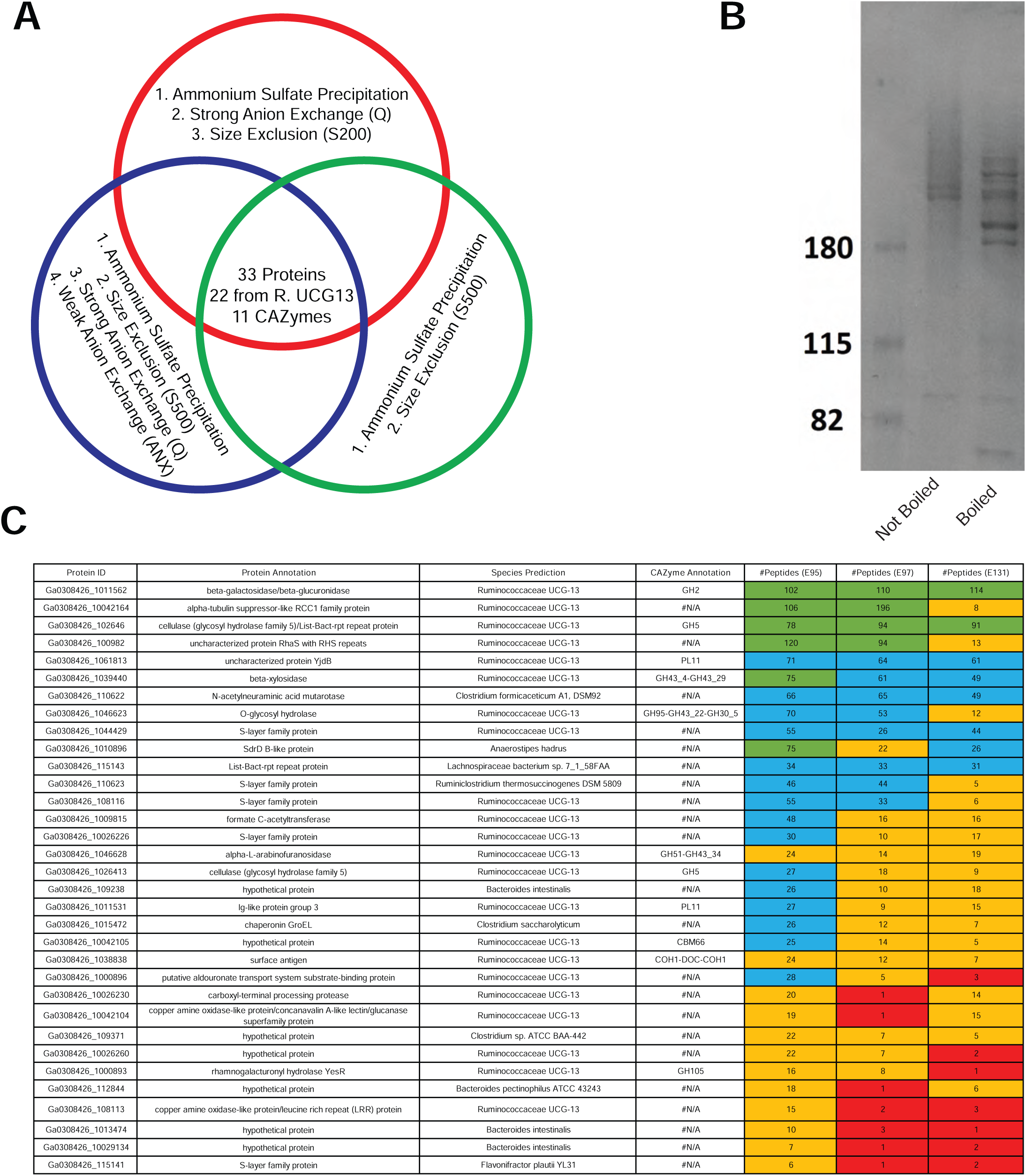
Activity-guided fractionation and proteomics narrow list of potential xanthanases. **a**, Venn diagram depicting activity-guided fractionation of culture supernatants followed by proteomic identification of candidate proteins present across all active preparations. **b**, SDS-PAGE of one of the fractionated culture supernatants (ANX processed sample) submitted for proteomic analysis (without and with boiling prior to analysis). Protein complexes or fragments that are larger are retained at the top of the gel while smaller proteins migrate towards the bottom of the gel. The ladder on the left shows how 180, 115, and 82 kDa standards are retained by the gel matrix. Boiling results in separation of protein complexes that cause streaking in the first lane and resolution into single protein bands that are denatured and migrate with respect to size. **c**, Proteomics narrows potential xanthanases to 33 proteins, 22 of which were from R. UCG13. Final candidates were obtained by collating proteins that were present across all three activity-based fractionation experiments with proteomic identification. Colors assist with visualizing number of peptides associated with each protein at different thresholds (<5, red; 5-24, orange; 25-74, blue; ≥75, green)

**Extended Data 6.**
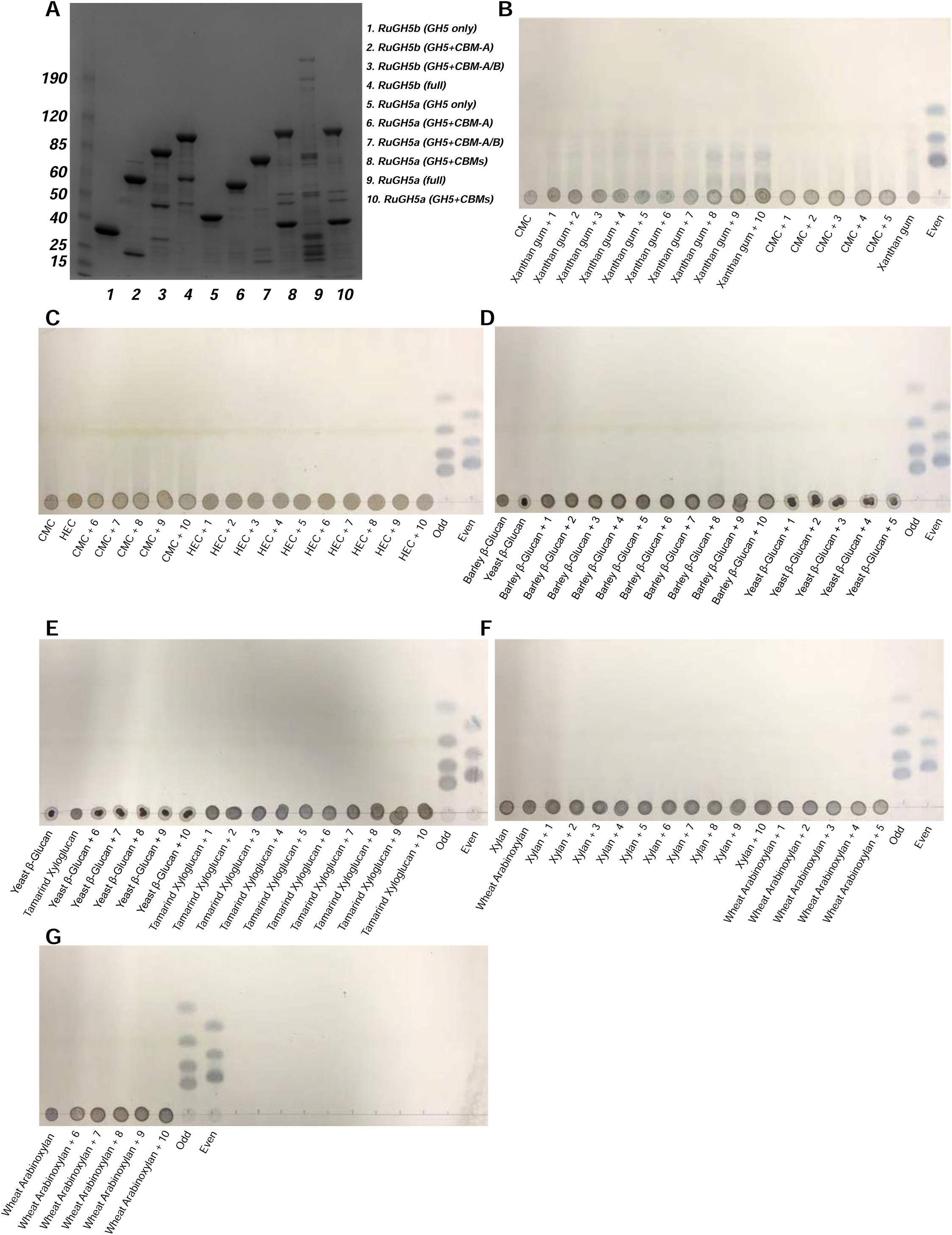
Activity of R. UCG13 GH5 enzymes on various polysaccharides. **a**, SDS-PAGE gel of purified GH5 constructs and their resultant activity as assessed by TLC on **b**, xanthan gum, **b-c**, carboxymethyl cellulose (CMC), **c**, hydroxyethyl cellulose (HEC), **d**, barley β-glucan, **d-e**, yeast β-glucan, **e**, tamarind xyloglucan, **f**, xylan, and **f-g**, wheat arabinoxylan. Enzymes are 1, *Ru*GH5b (GH5 only); 2, *Ru*GH5b (GH5 with CBM-A); 3, *Ru*GH5b (GH5 with CBM-A/B); 4, *Ru*GH5b (full protein); 5, *Ru*GH5a (GH5 only); 6, *Ru*GH5a (GH5 with CBM-A); 7, *Ru*GH5a (GH5 with CBM-A/B); 8, *Ru*GH5a (GH5 with CBM-A/B/C); 9, *Ru*GH5a (full protein); 10, replicate of 8. Strong color development in circles at baseline is undigested polysaccharide while bands or streaking that migrated with solvent are digested oligosaccharides and monosaccharides. Although minor streaking appears in some substrates due to residual oligosaccharides, comparing untreated substrate with enzyme incubated substrate allows determination of enzyme activity. *Ru*GH5a constructs with all 3 CBMs (8-10) showed clear activity on XG.

**Extended Data 7.**
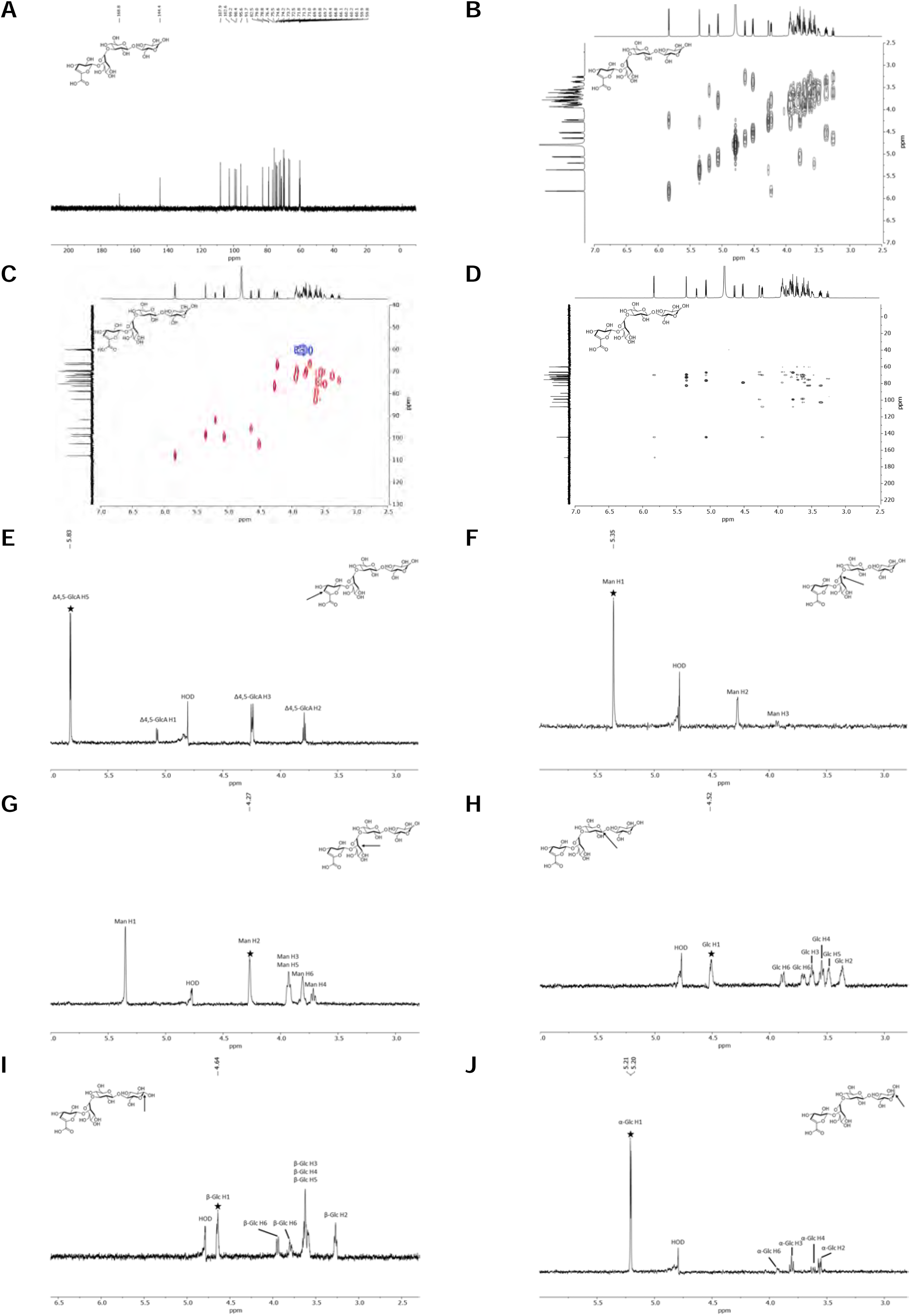
NMR of tetrasaccharide produced by *Ru*GH5a (and PL8 xanthan lyase). **a**, ^13^C-NMR spectrum of **tetrasaccharide** in D_2_O (^13^C: 150 MHz). **b**, COSY (^1^H-^1^H) spectrum of **tetrasaccharide** in D_2_O (600 MHz). **c**, HSQC (^1^H-^13^C) spectrum of **tetrasaccharide** in D_2_O. Red contours represent CH and CH_3_ groups, blue contours represent CH_2_ groups. **d**, HMBC (^1^H-^13^C) spectrum of **tetrasaccharide** in D_2_O. **e**, Selective 1H 1D-TOCSY spectrum of **tetrasaccharide** in D_2_O (^1^H: 600 MHz). Starred peak indicates the frequency irradiated (5.83 ppm) and arrow on the structure illustrates corresponding proton position being irradiated (Δ4,5-GlcA H5). **f**, Selective 1H 1D-TOCSY spectrum of **tetrasaccharide** in D_2_O (^1^H: 600 MHz). Starred peak indicates the frequency irradiated (5.35 ppm) and arrow on the structure illustrates corresponding proton position being irradiated (Man H1). **g**, Selective 1H 1D-TOCSY spectrum of **tetrasaccharide** in D_2_O (^1^H: 600 MHz). Starred peak indicates the frequency irradiated (4.27 ppm) and arrow on the structure illustrates corresponding proton position being irradiated (Man H2). **h**, Selective 1H 1D-TOCSY spectrum of **tetrasaccharide** in D_2_O (^1^H: 600 MHz). Starred peak indicates the frequency irradiated (4.52 ppm) and arrow on the structure illustrates corresponding proton position being irradiated (Glc H1). **i**, Selective 1H 1D-TOCSY spectrum of **tetrasaccharide** in D_2_O (^1^H: 600 MHz). Starred peak indicates the frequency irradiated (4.64 ppm) and arrow on the structure illustrates corresponding proton position being irradiated (β-Glc H1). **j**, Selective 1H 1D-TOCSY spectrum of **tetrasaccharide** in D_2_O (^1^H: 600 MHz). Starred peak indicates the frequency irradiated (5.20 ppm) and arrow on the structure illustrates corresponding proton position being irradiated (α-Glc H1). ^1^H-NMR analysis illustrated an inconsistent spectrum to that of the known degradation product from other xanthanases (including GH9 from *Paenibacillus nanensis* or the β-D-glucanase in *Bacillus* sp. Strain GL1), that hydrolyze xanthan at the reducing end of the branching glucose. LCMS analysis was consistent with a **tetrasaccharide** containing a Δ4,5-ene-GlcA moiety, but despite the appropriate mass, these differences in the ^1^H-NMR suggested an alternative cut site. To confirm, full structural elucidation was conducted by extensive NMR-analysis. In a similar fashion to Wilson and coworkers, spin systems for each monosaccharide were established via selective 1D-TOCSY experiments, selectively irradiating individual anomeric protons between *δ*_H_ 4.52 and *δ*_H_ 5.35, and the one vinylic proton of the Δ4,5-ene-GlcA residue at *δ*_H_ 5.83. This vinylic proton was easily identified by HSQC analysis via its distinct ^1^*J*_H,C_ correlation (*δ*_C_ 107.9 ppm), and its HMBC correlations to C-5 (*δ*_C_ 144.4 ppm) and C-6 (*δ_C_* 168.8 ppm) of Δ4,5-ene-GlcA. This proton was used as a starting point for structural elucidation, and in conjunction with the data obtained from 2D-HSQC and selective 1D-TOCSY experiments allowed for identification of the remainder of the Δ4,5-ene-GlcA spin system, including the anomeric position (*δ*_H_ 5.06, *δ*_C_ 99.2). Further HMBC analysis identified a correlation between H-1 of Δ4,5-ene-GlcA and the C-2 position of Man (*δ*_C_ 76.4 ppm). The inverse HMBC correlation was also observed from H-2 of the Man residue (*δ*_H_ 4.27) to the anomeric carbon of Δ4,5-ene-GlcA (*δ*_C_ 99.2), confirming the expected connectivity through a 1 → 2 linkage. COSY analysis identified a correlation between H-2 and H-1 (*δ*_H_ 5.35) of the of the α-Man residue, and as was the case for Δ4,5-ene-GlcA, the remainder of the positions were assigned from 2D-HSQC and selective 1D-TOCSY experiments. Interestingly, the anomeric position of α-Man appeared as a singlet, in contrast to the reported two anomeric proton signals associated with this residue in the tetrasaccharide isolated by Wilson. HMBC analysis from this anomeric position showed correlations to C-2, C-3, and C-5 (*δ*_C_ 76.4, *δ*_C_ 69.4, and *δ*_C_ 72.5 respectively) of Man. An additional correlation was observed to a carbon external to the Man subunit, with a chemical shift of 82.5 ppm. This shift was identified as belonging to the C-3 position of a nonreducing glucosyl residue. This was confirmed via HMBC correlations from both H-2 and H-4 of Glc(n.r). The H-2 peak was free of any overlap in the ^1^H-NMR spectrum, allowing for unambiguous assignment through a COSY correlation between itself and H-1 (*δ*_H_ 3.37 and *δ*_H_ 4.52 respectively), as well as the H-3 proton with a chemical shift of 3.63 ppm. This gave us confidence that the α-Man residue was connected via a 1 → 3 linkage to this Glc(n.r) subunit. Importantly, the anomeric proton of the Glc(n.r) residue appeared as a single doublet, integrating with a value of one in the ^1^H-NMR spectrum with a coupling constant of 8.0 Hz, consistent with a β-configured Glc(n.r) monomer. This confirmed connectivity between the α-Man and β-Glc(n.r) residues, suggesting a disparate structure to the previously reported degradation product. Finally, a key HMBC correlation was observed from the anomeric proton (*δ*_H_ 4.52) to an external carbon with a ^13^C-chemical shift of ∼79 ppm. Upon closer inspection, this carbon was actually two separate peaks, corresponding to the C-4 position of the alpha (minor) and beta (major) anomers (*δ*_C_ 79.0 and *δ*_C_ 78.8 respectively) of the reducing Glc. This confirmed the expected 1 → 4 linkage of the backbone Glc residues, albeit illustrating hydrolysis had occurred at the reducing end of the nonbranching Glc. To complete structural elucidation, the remainder of the positions were assigned from 2D-HSQC and selective 1D-TOCSY experiments for both the alpha and beta anomers separately.

**Extended Data 8.**
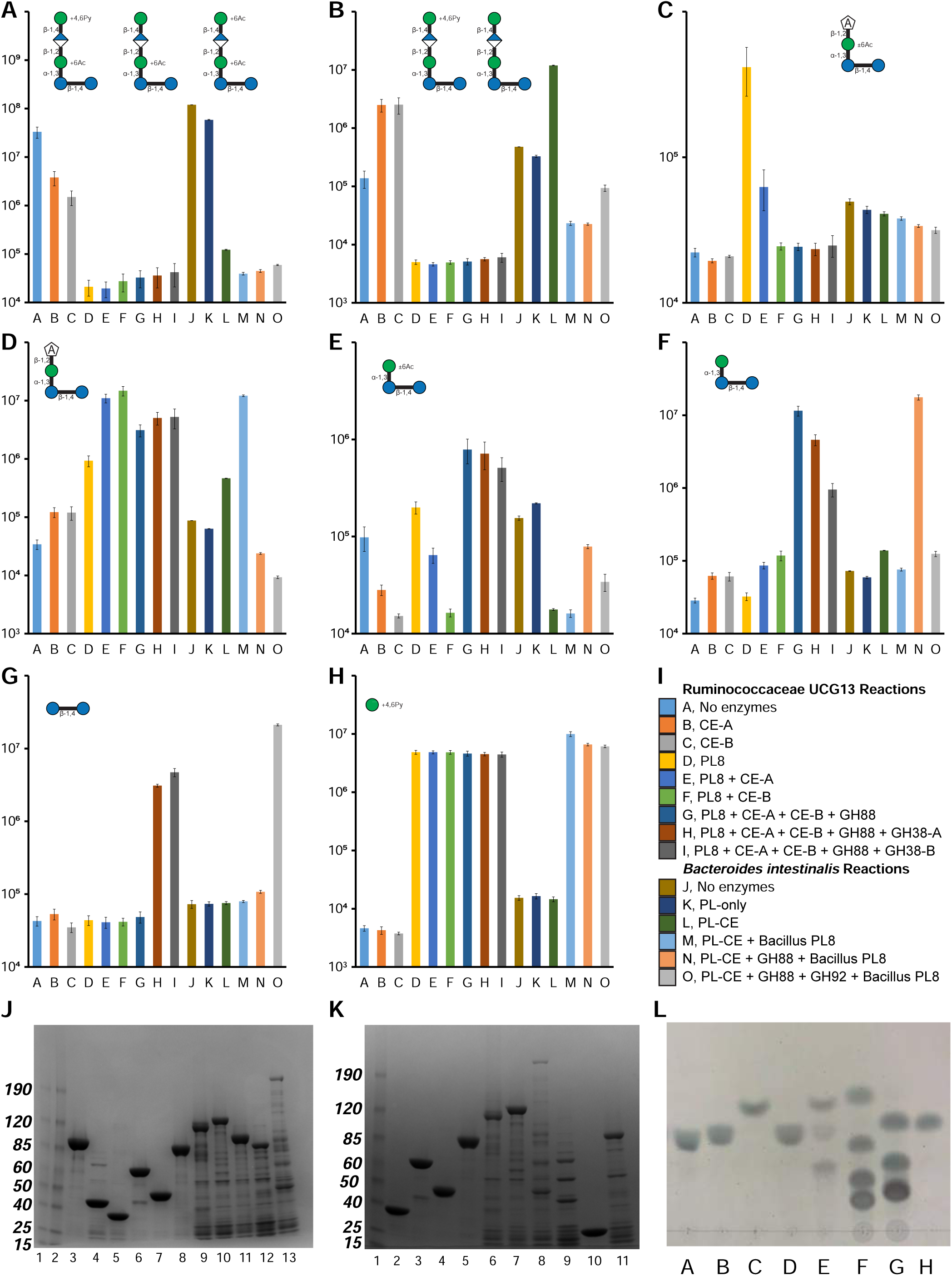
Activity of R. UCG13 and *B. intestinalis* enzymes. LC-MS analysis was used to track relative increases and decreases of intermediate oligosaccharides with the addition of enzymes, verifying their abilities to successively cleave XG pentasaccharides to their substituent monosaccharides. Integrated extracted ion counts (n=4, SEM) that correlate with compound abundance are shown for **a**, acetylated pentasaccharide (M-H ions: 883.26, 953.26, 925.27), **b**, deacetylated pentasaccharide (M-H ions: 841.25, 911.25), **c**, acetylated tetrasaccharide (2M-H ion: 1407.39), **d**, tetrasaccharide (M-H ion: 661.18), **e**, acetylated trisaccharide (M+Cl ion: 581.15), **f**, trisaccharide (M+Cl ion: 539.14), **g**, cellobiose (M+Cl ion: 377.09), and **h**, pyruvylated mannose (M-H ion: 249.06). Reactions were carried out using xanthan oligosaccharides produced by the *Ru*GH5a to test activities of the R. UCG13 (A-I) and *B. intestinalis* (J-O) enzymes. R. UCG13 enzymes were tested in reactions that included (A) no enzyme, (B) R. UCG13 CE-A, (C) R. UCG13 CE-B, (D) R. UCG13 PL8, (E) R. UCG13 PL8 and CE-A, (F) R. UCG13 PL8 and CE-B, (G) R. UCG13 PL8, both CEs, and GH88, (H) R. UCG13 PL8, both CEs, GH88, and GH38-A, (I) R. UCG13 PL8, both CEs, GH88, and GH38-B. *B. intestinalis* enzymes were tested in reactions that included (J) no enzyme, (K) Bi PL-only, (L) Bi PL-CE, (M) Bi PL-CE and Bacillus PL8, (N) Bi PL-CE and GH88 and Bacillus PL8, (O) Bi PL-CE, GH88, and GH92 and Bacillus PL8. **i**, Legend of enzymes included in each reaction. Recombinant enzymes were purified and analyzed for expression and purity by SDS-PAGE. Proteins generally expressed well with a single dense band indicating overexpression of the enzyme at its predicted molecular weight as compared to a size ladder. Exceptions included the R. UCG13 GH88 and CE-A, both of which had bands at the predicted enzyme size but also showed bands of comparable density at other sizes resulting from either proteolysis or co-purification of undesired *E. coli* proteins. **j**, SDS-PAGE gel of purified enzymes with 4.5 µg loaded, including (1-2) ladder, (3) *B. intestinalis* GH3, (4) *B. intestinalis* GH5, (5) *B. intestinalis* PL-only, (6) *B. intestinalis* PL-CE, (7) *B. intestinalis* GH88, (8) *B. intestinalis* GH92, (9) R. UCG 13 GH38-A, (10) R. UCG13 GH38-B, (11) R. UCG13 GH94, (12) R. UCG13 PL8, (13) R. UCG13 CE-A. **k**, SDS-PAGE gel of purified enzymes with 4.5 µg loaded, including (1) ladder, (2) *B. intestinalis* PL-only, (3) *B. intestinalis* PL-CE, (4) *B. intestinalis* GH88, (5) *B. intestinalis* GH92, (6) R. UCG13 GH38-A, (7) R. UCG13 GH38-B, (8) R. UCG13 CE-A, (9) R. UCG13 GH88, (10) R. UCG13 CE-B, (11) R. UCG13 PL8. **l**, TLC analysis showed that R. UCG13 GH94 and *B. intestinalis* GH3 are active on cellobiose. From left to right lane show (A) *Ru*GH5b (full protein), (B) *Ru*GH5a (full protein), (C) *B. intestinalis* GH3, (D) *B. intestinalis* GH5, (E) R. UCG13 GH94, (F) odd standards, (G) even standards, (H) cellobiose. Odd and even standards are maltooligosaccharides with 1, 3, 5, and 7 hexoses or 2, 4, and 6 hexoses, respectively. While the *B. intestinalis* GH3 only produced one product, the R. UCG13 GH94 produced two, one matching the approximate Rf of glucose while the other had a much lower Rf which presumably is phosphorylated glucose (matching the known phosphorylase activity of the GH94 family).

**Extended Data 9.**
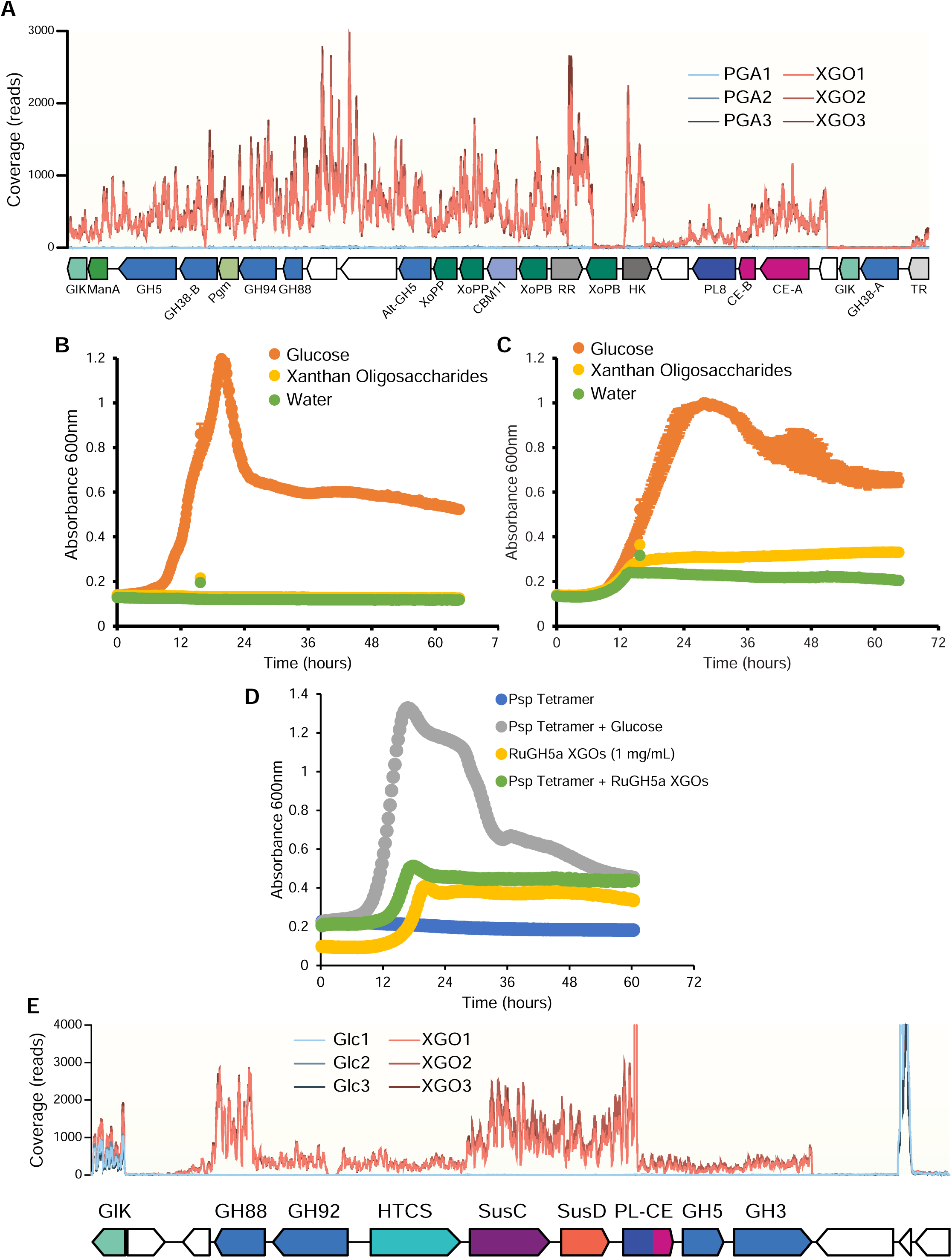
**a**, Traces of RNA-seq expression data from triplicates of the original culture grown on either XG or polygalacturonic acid (PGA), illustrating overexpression of the XG PUL. **b**, *Bacteroides clarus* and **c**, *Parabacteroides distasonis* isolated from the original culture did not grow on XGOs. **d**, *Bacteroides intestinalis* did not grow on tetramer generated with *P. nanensis* GH9 and PL8 (Psp Tetramer) even in the presence of 1 mg/mL *Ru*GH5a generated XGOs to activate the PUL. Growth on glucose confirmed that the Psp Tetramer was not inherently toxic to cells. All substrates were used at 5 mg/mL unless otherwise noted. Growths are n ≥ 2, error bars show SEM (in most cases, smaller than the marker). **e,** Traces of RNA-seq expression data from triplicates of *B. intestinalis* grown on either glucose (Glc) or XG oligosaccharides (XGOs), illustrating overexpression of the XGO PUL.

**Extended Data 10.**
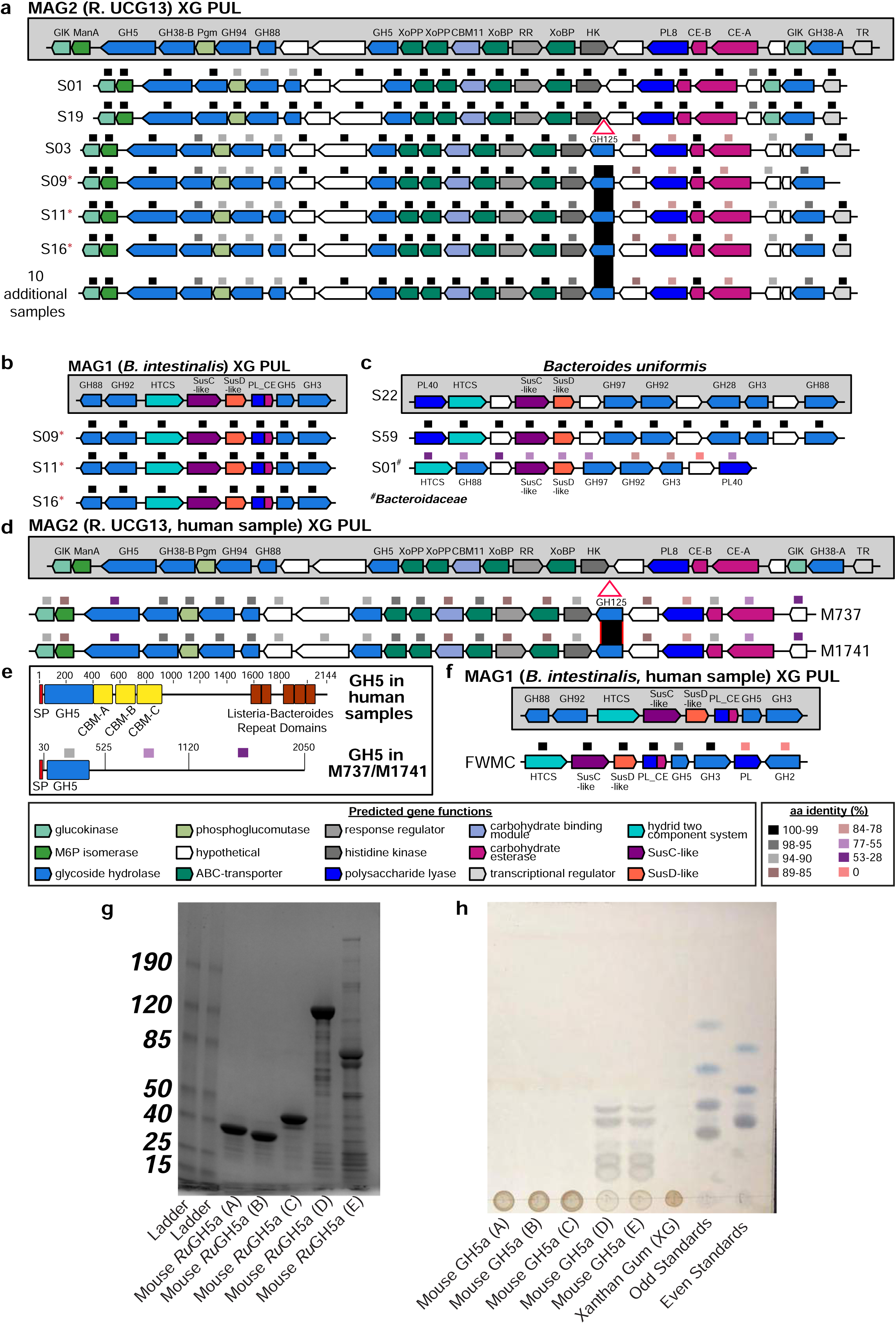
**a,** Metagenomic sequencing of additional 16 cultures (S, human fecal sample) that actively grew on and degraded xanthan gum revealed two architectures of the R. UCG13. The more prevalent locus contained a GH125 insertion. The 10 additional samples with this locus architecture include: S22, S25, S39, S43, S44, S45, S49, S53, S58, and S59. **b,** The *B. intestinalis* xanthan locus was present in 3 additional cultures. **c,** Additional members of the Bacteroideceae family harbor a PUL with a GH88, GH92 and GH3 that could potentially enable utilization of XG-oligosaccharides. **d,** The GH125-containing version of the R. UCG13 xanthan locus was detected in two mouse fecal samples (M, mouse fecal sample). **e,** Comparison of the human and mouse *Ru*GH5a aa sequence, showing the annotated signal peptide (SP), GH5 domain, three carbohydrate binding modules (CBMs), and multiple Listeria-Bacteroides repeat domains. **f,** Genetic organization and aa identity (%) between the *B. intestinalis* xanthan locus in the original human sample and a PUL detected in a fracking water microbial community (FWMC) using LAST-searches. **g**, SDS-PAGE gel of purified enzymes with 4.5 µg loaded, including ladder and the different mouse *Ru*GH5a constructs. A, B, and C are all versions of the GH5 domain alone, D is a construct designed to terminate at a site homologous to the last CBM in the human *Ru*GH5a, and E is a full-length construct of the mouse *Ru*GH5a. **h**, TLC of each mouse *Ru*GH5a construct incubated with XG and also odd (1, 3, 5, and 7 residues) and even (2, 4, and 6 residues) malto-oligosaccharide standards. The GH5-only constructs did not degrade XG, but constructs D and E (with regions homologous to the human *Ru*GH5a CBMs) were able to hydrolyze XG.

